# The wheat blast pathogen *Pyricularia graminis-tritici* has complex origins and a disease cycle spanning multiple grass hosts

**DOI:** 10.1101/203455

**Authors:** Vanina L. Castroagudín, Anderson L. D. Danelli, Silvino I. Moreira, Juliana T. A. Reges, Giselle de Carvalho, João L.N. Maciel, Ana L. V. Bonato, Carlos A. Forcelini, Eduardo Alves, Bruce A. McDonald, Daniel Croll, Paulo C. Ceresini

**Author notes:** These authors contributed equally to this manuscript (Vanina L. Castroagudín, Anderson L. D. Danelli, Silvino I. Moreira, Juliana T. A. Reges).

## Abstract

The wheat blast disease has been a serious constraint for wheat production in Latin America since the late 1980s. We used a population genomics analysis including 95 genome sequences of the wheat blast pathogen *Pyricularia graminis-tritici* (*Pygt*) and other *Pyricularia* species to show that *Pygt* is a distinct, highly diverse pathogen species with a broad host range. We assayed 11 neutral SSR loci in 526 *Pygt* isolates sampled from wheat and other grasses distributed across the wheat-growing region of Brazil to estimate gene flow, assess the importance of sexual reproduction, and compare the genetic structures of *Pygt* populations infecting wheat and nearby grasses. Our results suggest a mixed reproductive system that includes sexual recombination as well as high levels of gene flow among regions, including evidence for higher gene flow from grass-infecting populations and into wheat-infecting populations than vice versa. The most common virulence groups were shared between the grass- and wheat-infecting *Pygt* populations, providing additional evidence for movement of *Pygt* between wheat fields and nearby grasses. Analyses of fruiting body formation found that proto-perithecia and perithecia developed on senescing stems of wheat and other grass hosts, suggesting that sexual reproduction occurs mainly during the saprotrophic phase of the disease cycle on dead residues. *Phalaris canariensis* (canarygrass) supported the fullest development of perithecia, suggesting it is a promising candidate for identifying the teleomorph in the field. Based on these findings, we formulated a more detailed disease cycle for wheat blast that includes an important role for grasses growing near wheat fields. Our findings strongly suggest that widely grown pasture grasses function as a major reservoir of wheat blast inoculum and provide a temporal and spatial bridge that connects wheat fields across Brazil.

**Author summary:** After the first wheat blast epidemic occurred in 1985 in Paraná, Brazil, the disease spread to Bolivia, Argentina, and Paraguay, and was introduced into Bangladesh in 2016 followed by India in 2017. Wheat blast is caused by *Pyricularia graminis-tritici* (*Pygt*), a highly diverse pathogen species related to the rice blast fungus *P. oryzae*, but with an independent origin and a broader host range. We conducted a large scale contemporary sampling of *Pygt* from symptomatic wheat and other grass species across Brazil and analyzed the genetic structure of *Pygt* populations. *Pygt* populations on both wheat and other grasses had high genotypic and virulence diversity, a genetic structure consistent with a mixed reproductive system that includes regular cycles of recombination. The pathogen formed sexual fruiting structures (perithecia) on senescing stems of wheat and other grasses. Historical migration analyses indicated that the majority of gene flow has been from *Pygt* populations on other grasses and into the *Pygt* population infecting wheat, consistent with the hypothesis that *Pygt* originated on other grasses before becoming a wheat pathogen. We found that the *Pygt* populations infecting wheat were indistinguishable from the *Pygt* populations infecting other grass species, including signal grass (*Urochloa brizantha*). Because *U. brizantha* is a widely grown grass pasture often found next to wheat fields, we propose that it functions as reservoir of *Pygt* inoculum that provides a temporal and spatial bridge that connects wheat fields in Brazil.

## Introduction

*Pyricularia* is a species-rich genus including many fungal pathogens that show specialization towards different host species in the Poaceae family, including rice (*Oryza sativa*), wheat (*Triticum aestivum*), oat (*Avena sativa*), barley *(Hordeum vulgare*), and millets (*Eleusine coracana, Pennisetum glaucum*, *Setaria italica*), as well as more than 50 other species of grasses [1–5]. Several studies indicated that distinct *Pyricularia* species emerged through repeated radiation events from a common ancestor [6, 7]. Such radiation events often result from ecological adaptations that include host jumps or shifts and changes in pathogenicity [4, 8]. These ecological adaptations may lead to the emergence of new species of "domesticated" host-specialized fungal pathogens infecting agricultural crops from "wild" ancestral source populations found on undomesticated plants [4, 8]. Examples of speciation following host specialization are common in cereal agro-ecosystems and were already described for several plant pathogenic fungi, including *Pyricularia oryzae* on rice and *P*. *grisea* on *Digitaria* spp. [5], *Zymoseptoria tritici* on wheat [9], *Rhynchosporium commune* on barley [10], *Ceratocystis fimbriata* on cacao (*Theobroma cacao*), sweet potato (*Ipomoea batatas*) and sycamore (*Platanus* spp.) [11], and *Microbotryum violaceum* on *Silene* spp. [12]. For *P*. *oryzae*, causal agent of rice blast [5, 13], strains that infect rice are thought to have emerged by ecological adaptation via host shifts from millet (*Setaria* spp.) to rice and to have co-evolved with their respective hosts during the domestication of rice and millet in China about 7000 BC [14].

A previous study indicated that a new *Pyricularia* species, named *Pyricularia graminis-tritici* (*Pygt*), emerged in southern Brazil during the last century as the pathogen causing wheat blast [15]. *Pygt* is closely related to *P. oryzae* [15]. Wheat blast was first reported in Paraná State, Brazil in 1985 [16, 17] and since then has become an increasingly important disease, causing crop losses ranging from 40% to 100% [18]. Blast disease has also been reported in other important crops growing in the same agro-ecosystems in Latin America, including pastures of signal grass (*Urochloa brizantha, ex Brachiaria brizantha*), barley, oats, rye (*Secale cereale*), and triticale (*x Triticosecale*). Although other *Pyricularia* species can cause blast symptoms on wheat, we focused this study on *Pygt*, which is the major species associated with wheat blast [15, 17, 19–24]. Since its discovery, *Pygt* has spread across all wheat-cropping areas in Brazil [17, 18, 25–27] and is now found in Bolivia, Argentina and Paraguay [28]. Its first report outside South America was an outbreak in Bangladesh in 2016 [29–31] followed by its spread to India in 2017 [32, 33]. Wheat blast is a major quarantine disease in the United States [27] and it is considered a threat to wheat cultivation in disease-free areas across Asia, Europe, and North America [34].

*Pygt* can be dispersed over short and long distances by aerial inoculum (conidia) [35] and also on infected seeds [36]. Unlike most *Pyricularia* species, *Pygt* isolates recovered from wheat can infect a wide range of hosts, including the tribes *Hordeae, Festuceae, Avenae, Chlorideae, Agrosteae* and *Paniceae* [37]. Under natural field conditions, close physical proximity between cultivated plants and other poaceous hosts (i.e., weeds or invasive grass species) could enable genetic exchange among *Pyricularia* populations on different hosts and facilitate host shifts. Cross-infection and inter-fertility between fungal strains from different grass hosts were hypothesized to play a role in the emergence of wheat blast [38, 39]. Evidence to support this hypothesis was presented in a recent study that analyzed variation in the avirulence genes *PWT3* and *PWT4* [40]. This study proposed that wheat blast emerged via a host shift from a *Pyricularia* population infecting *Lolium*. In their model, a *Lolium*-derived isolate carrying the Ao avirulence allele at the *PWT3* locus infected a susceptible wheat cultivar carrying the *rwt3* susceptibility allele. The model further proposes that the spread of wheat blast in the 1980s was enabled by the widespread cultivation in Brazil of susceptible wheat cultivars carrying *rwt3*. Selection on less common *Rwt3* wheat cultivars favored the emergence of pathogen strains with non-functional *PWT3* alleles, and the authors proposed that it was these *pwt3* strains that eventually became the epidemic wheat blast population found in South America.

*Pyricularia* is considered a genus of pathogens with high evolutionary potential [39, 41, 42]. The evolutionary potential of a pathogen population reflects its ecology and biology, and its population genetic structure [41, 42]. Pioneering studies on the genetic structure of *Pygt* indicated a highly variable population distributed across different Brazilian states [43, 44]. Analyses of three regional populations sampled in Brazil between 2005 and 2008 suggested long distance gene flow and a mixed reproductive system [39]. These findings indicated that *Pygt* is a pathogen with high evolutionary potential, according to the risk model proposed by McDonald and Linde [41, 42].

Knowledge about the evolutionary potential of *Pygt* populations is needed to predict the durability of genetic resistance to wheat blast. An intense search for blast resistance began with the first report of the disease more than 30 years ago, but breeding success has been erratic and inconsistent [45–48]. The average durability of resistant wheat varieties has been only two to three years [49]. Furthermore, wheat genotypes behaved differently in different regions, indicating genotype-by-environment interactions or a region-specific distribution of virulence groups [50]. Given that *Pygt* is now present in all Brazilian wheat growing areas [15, 28], it is likely that both the incidence and severity of wheat blast are affected by the virulence groups that predominate in each region [39]. In fact, the occurrence of virulence groups in *Pygt* populations was already described [39, 43, 50, 51], but information about the virulence composition and genetic structure of contemporary populations of the wheat blast pathogen remains limited.

Several lines of evidence indicate that *Pygt* populations recombine regularly in Brazil: both mating types and fertile strains were present in wheat fields, field populations contain high genetic diversity, and gametic equilibrium is found among neutral marker loci [26, 39, 52]. Under laboratory conditions, *Pygt* isolates showed the capacity for sexual reproduction [37] and were shown to be sexually compatible with *Pyricularia* isolates from other poaceous hosts including plantain signalgrass (*Urochloa plantaginea, ex Brachiaria plantaginea*), goosegrass (*Eleusine indica*), finger-millet (*Setaria italica*), rescuegrass (*Bromus catharticus*), canary grass (*Phalaris canariensis*) and triticale (x *Triticosecale*) [52, 53]. Crosses between isolates recovered from wheat and *Urochloa plantaginea* produced perithecia with asci and ascospores, a clear indicator of sexual reproduction [54], but perithecia have not yet been found in blasted wheat fields and it remains unclear where and when the sexual stage occurs.

Here we bring together findings from a series of experiments conducted to better understand the origins of wheat blast and formulate an improved disease cycle. We first used population genomic analyses including 36 *Pygt* strains originating from many different hosts and 59 strains of other *Pyricularia* species to infer the genealogical relationships among *Pyricularia* species and better define the phylogenetic boundaries of *Pygt*. We next generated and analyzed a microsatellite dataset from 526 contemporary Brazilian isolates of *Pygt* sampled from wheat fields and invasive grasses across Brazil to compare the genetic structures of *Pygt* populations found on wheat and other grasses. We then compared the distribution of *Pygt* virulence groups found in wheat fields with the distribution of virulence groups found on invasive grasses growing in or near those wheat fields. Finally, we conducted experiments to identify grass hosts and tissues where sexual perithecia are most likely to form to better understand the importance of sexual recombination in *Pygt* population biology and identify the hosts most likely to support formation of the teleomorph. This combination of experiments provided novel insights into the origins and epidemiology of wheat blast.

## Results

### Several *Pyricularia* species were recovered from blast lesions on wheat and invasive grasses

We sampled *Pyricularia spp*. from wheat and other poaceous hosts in naturally infected wheat fields distributed across the seven states where wheat is grown in Brazil. Amongst the 556 *Pyricularia* spp. isolates included in our analyses, 30 isolates were not *Pygt* (Table 1, Supplementary Table 2). Based on the sequence of the hydrophobin *MPG1*, an isolate from DF-GOw was classified as *P. urashimae*. This was the only isolate recovered from a wheat head that was not *Pygt*. The 23 isolates from MSp included two isolates of *P. grisea* (recovered from *Digitaria sanguinalis*), 13 isolates of *P. pennisetigena* (from *Cenchrus echinatus, Eragrostis plana, Panicum maximum* and *Urochloa brizantha*), five isolates of *P. urashimae* (from *Avena sativa, Echinochloa crusgalli, P. maximum*, and *U. brizantha*), and three *Pyricularia* isolates that could not be identified at the species level (from *P. maximum* and *U. brizantha*). The five isolates found in PR_p_ included two isolates of *P. grisea* (from *D. sanguinalis*), one of *P. pennisetigena* (from *U. brizantha*), and two of *P. urashimae* (from *Chloris distichophylla* and *P. maximum*). Isolate 363 came from a rice field, probably from a *Digitaria* spp., and was classified as *P. grisea*.

**Table 1.** Populations of the blast pathogen *Pyricularia graminis-tritici* from wheat and other poaceous hosts characterized in this study.

| *Species, host, cultivar* | Population | Sampling Location, State | Coordinates | Sampling year | *N* |
| --- | --- | --- | --- | --- | --- |
| *Triticum aestivum* |  |  |  |  |  |
| Several cultivars | 2005<sub>w</sub> | Central-southern Brazil | ... | 2005 | 79 |
| BRS 254, BR18 | DF-GO<sub>w</sub> | Brasília, DF; and Rio Verde, GO | 17°19'46.8"S, 50°06'17.5"W | 2012-2013 | 86 |
| BRS 264, BR18 | MG<sub>w</sub> | Patrocínio, and Perdizes, MG | 19°09'10.1"S, 47°16'5.7"W | 2012-2013 | 62 |
| BRS Guamirim | MS<sub>w</sub> | Amambaí, and Aral Moreira, MS | 22°59'33.1"S, 55°20'11.1"W | 2012-2013 | 82 |
| CD 104 | PR<sub>w</sub> | Londrina, Jandaia do Sul, PR | 23°14'16.0"S, 51°10'10.5"W | 2012-2013 | 74 |
| ... | RS<sub>w</sub> | Passo Fundo, São Luiz Gonzaga, São Borja and Três de Maio, RS | 28°33'16"S, 55°21'52.5"W | 2012-2013 | 52 |
| CD 116 | SP<sub>w</sub> | Itaí, SP | 23°33'8"S, 49°3'24.9"W | 2012-2013 | 31 |
| **Total (*T. aestivum*)** |  |  |  |  | **466** |
| Other poaceous hosts |  |  |  |  |  |
| *Avena sativa*, *Cenchrus echinatus*, *Chloris distichophylla*, *Cynodon* spp., *Digitaria insularis*, *Digitaria sanguinalis*, *Echinochloa crusgalli*, *Eleusine indica*, *Eragrostis plana*, *Panicum maximum*, *Rhynchelytrum repens*, *Sorghum sudanense*, and *Urochloa brizantha*. | MS<sub>P</sub> | Amambaí and Aral Moreira, MS | 22°59'33.1"S, 55°20'11.1"W | 2012-2013 | 29 |
|  | PR<sub>P</sub> | Londrina, PR | 23°14'16.0"S, 51°10'10.5"W | 2012-2013 | 31 |
| **Total (Other poaceous hosts)** |  |  |  |  | **60** |
| **Total** |  |  |  |  | **526** |

### Population genomic analyses reveal that *Pyricularia graminis-tritici* comprises a single highly diverse species

Our first goal in this study was to infer the genealogical relationships among the *Pyricularia* species found in Brazil and to determine if the *Pygt* strains associated with blast on wheat and other grasses comprise a single species. We extended the analysis from Islam et al. [31] by adding into the genealogy Pyricularia isolates from 10 non-wheat hosts sampled in sympatry with 22 wheat blast isolates. The 47 *P. oryzae* strains associated with rice blast grouped together as a near-clonal genotype that was distinct from the group of 32 *Pygt* strains found on wheat and other grasses in Brazil and Bangladesh (Fig 1). The inferred genealogical relationships indicated that the *Pygt* strains sampled mainly from wheat comprise a single highly diverse species. The formerly described *P. oryzae* pathotype *Triticum* clade (indicated as *PoT* in the genealogy) [15] was not distinct from the *P. graminis-tritici* (*Pygt*) clade (Fig 1). The clade formed by *Pygt* strains sampled from infected wheat ears and other grass hosts contained much more polymorphism than the rice-infecting *P. oryzae* strains available in public genome databases. Despite the higher overall diversity, several of the Brazilian *Pygt* strains formed sub-clades that may represent expanded clonal lineages (Fig 1). In two of these sub-clades, closely related strains from the same sub-clade were found infecting different hosts.

**Fig 1.**
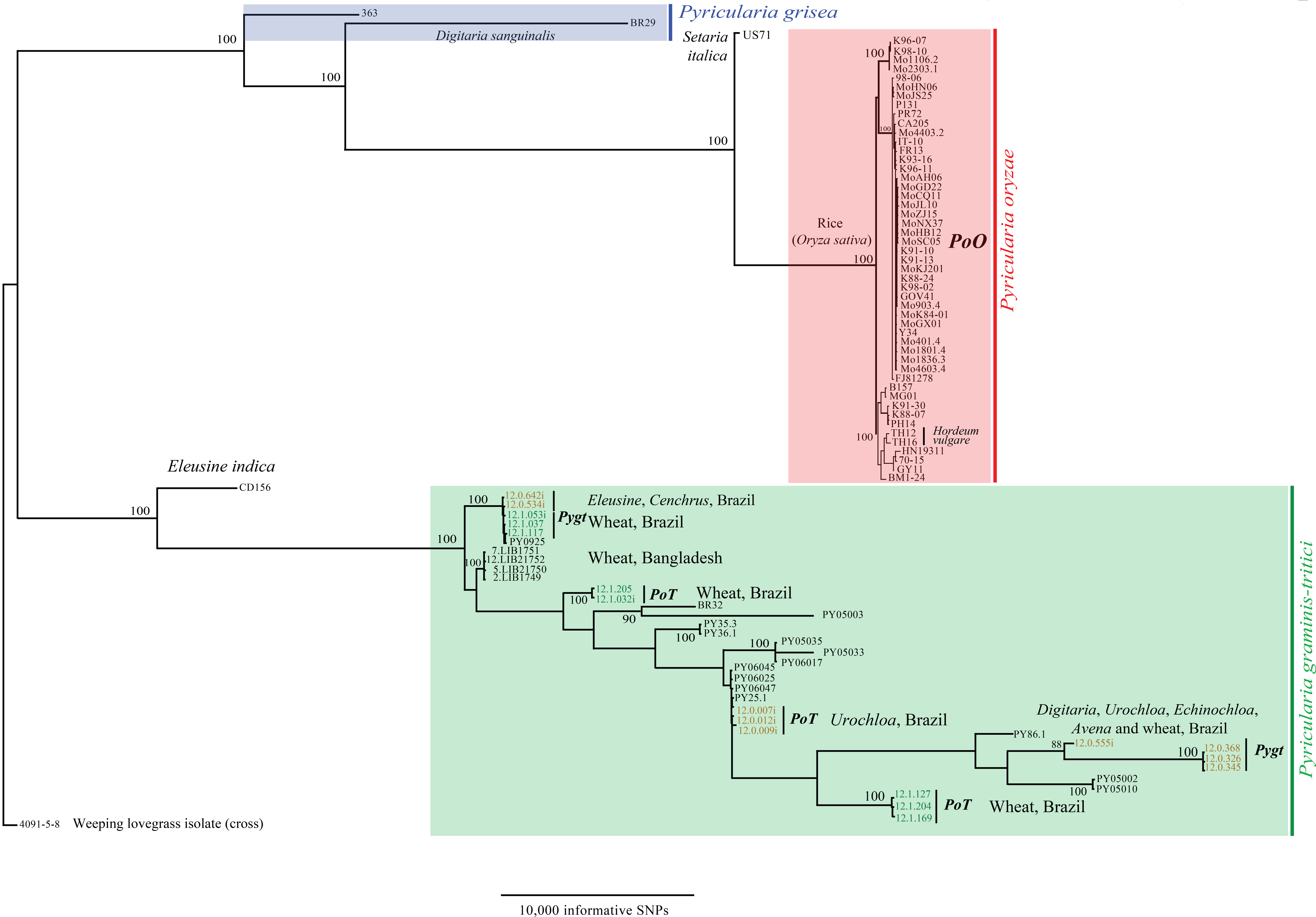
Population genomic analyses of transcriptomic single nucleotide polymorphisms among isolates of *Pyricularia graminis-tritici* from wheat and several other poaceous hosts in Brazil, *P. oryzae*, *P. grisea* from *Digitaria sanguinalis* and other *Pyricularia* spp. from *Setaria italica*, and *Eleusine indica*. The scale bar shows the number of informative sites. The samples included 47 rice blast strains with publically available genome sequences, 32 Brazilian wheat and other poaceous blast strains, seven strains from various additional hosts and four wheat blast samples collected in Bangladesh in spring 2016. The dataset contained only SNPs reliably called in the transcriptomic sequences of the Bangladesh sample 12 and genotyped in at least 90% of all other strains. We retained 55,041 informative SNPs. A maximum likelihood phylogeny was constructed using RAxML version 8.2.8 with a GTR substitution matrix and 100 bootstrap replicates. Pygt and PoT stands for the formerly described *P. graminis-tritici* and *P. oryzae* pathotype *Triticum*.

### Populations of *Pygt* from wheat and other grasses share genotypes

To explore the possibility of gene and genotype flow among the *Pygt* populations infecting wheat and other grasses, we conducted population genetic analyses using 11 neutral microsatellite (SSR) markers in an expanded dataset including 526 Brazilian *Pygt* isolates. A total of 198 different multilocus microsatellite genotypes (MLMGs) were found among the 526 isolates (Table 2, Fig 2). Of these MLMGs, 165 (83%) were found in only one population (Tables 2–4), but 33 MLMGs (17%) were shared by sympatric (from the same region) or allopatric (from different regions) populations of *Pygt*. These 33 MLMGs encompassed 257 isolates (224 from wheat, and 33 from other grasses), with 20 of these MLMGs (corresponding to 176 isolates) found exclusively on wheat. The number of MLMGs within a population that were shared across populations ranged from four (7 isolates) in SP_w_ to 15 (46 isolates) in MS. No MLMGs were shared between the isolates collected in 2005 and those collected in 2012 (Tables 3 and 4), indicating that *Pygt* clones do not persist over time.

**Table 2.**
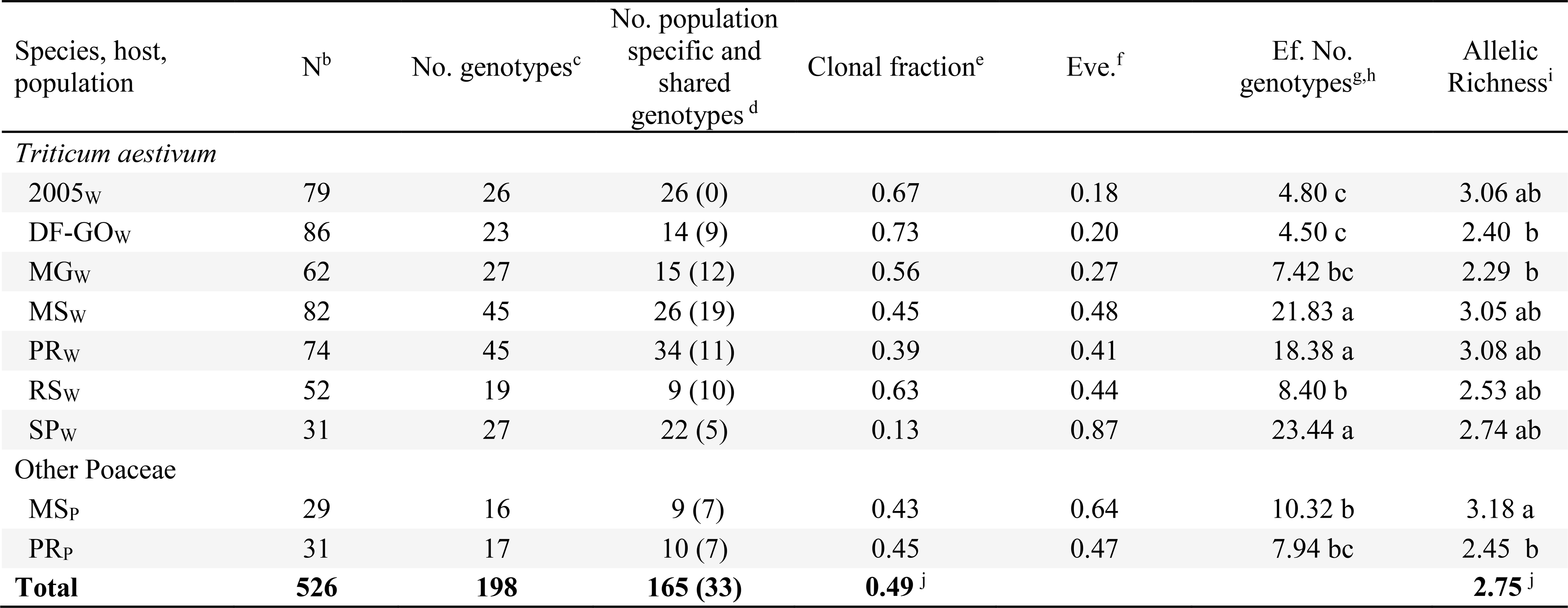
Measures of gene and genotypic and clonal diversity in populations of *Pyricularia graminis-tritici* from wheat and other poaceous hosts in Brazil^a^

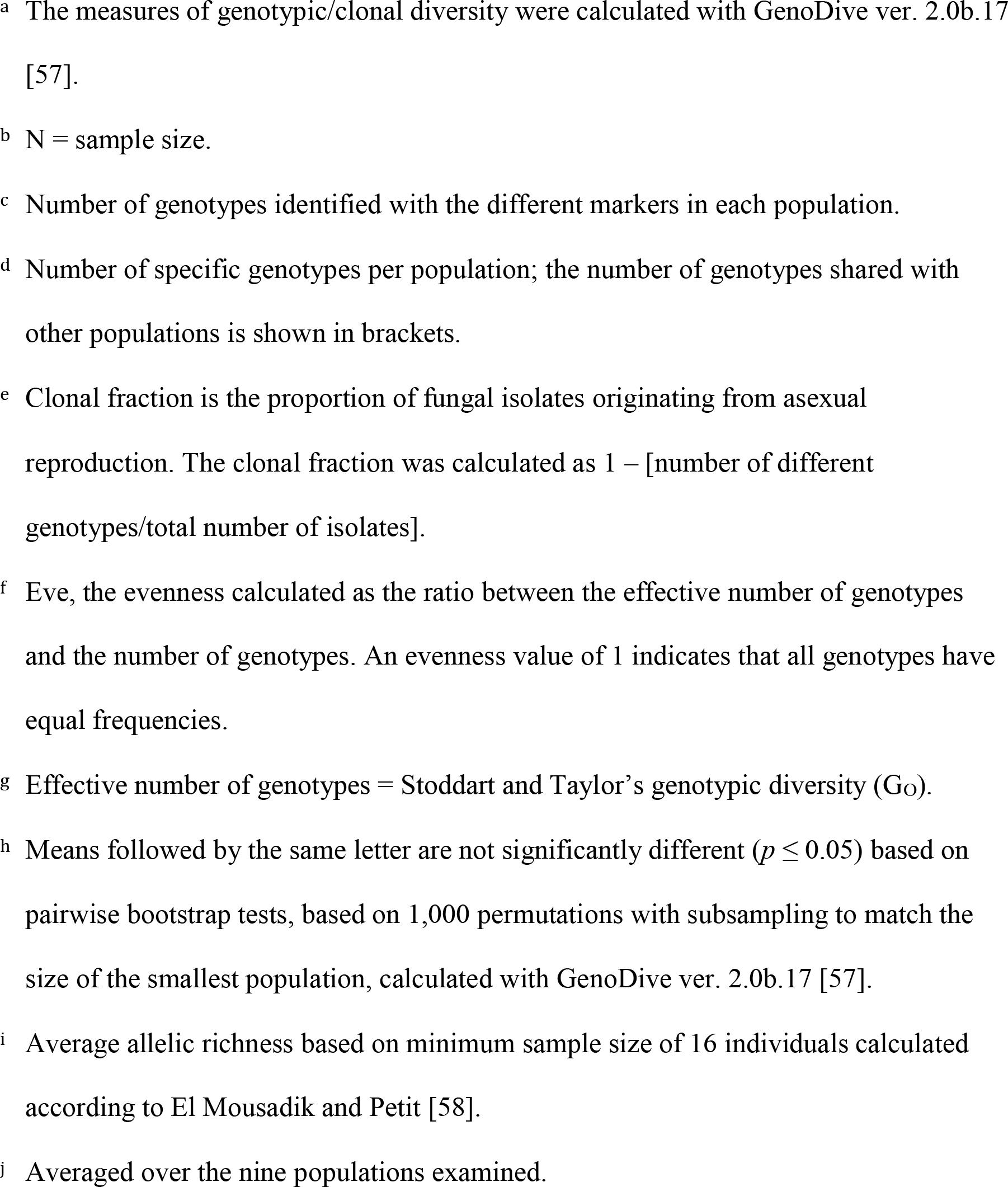

**Fig 2.**
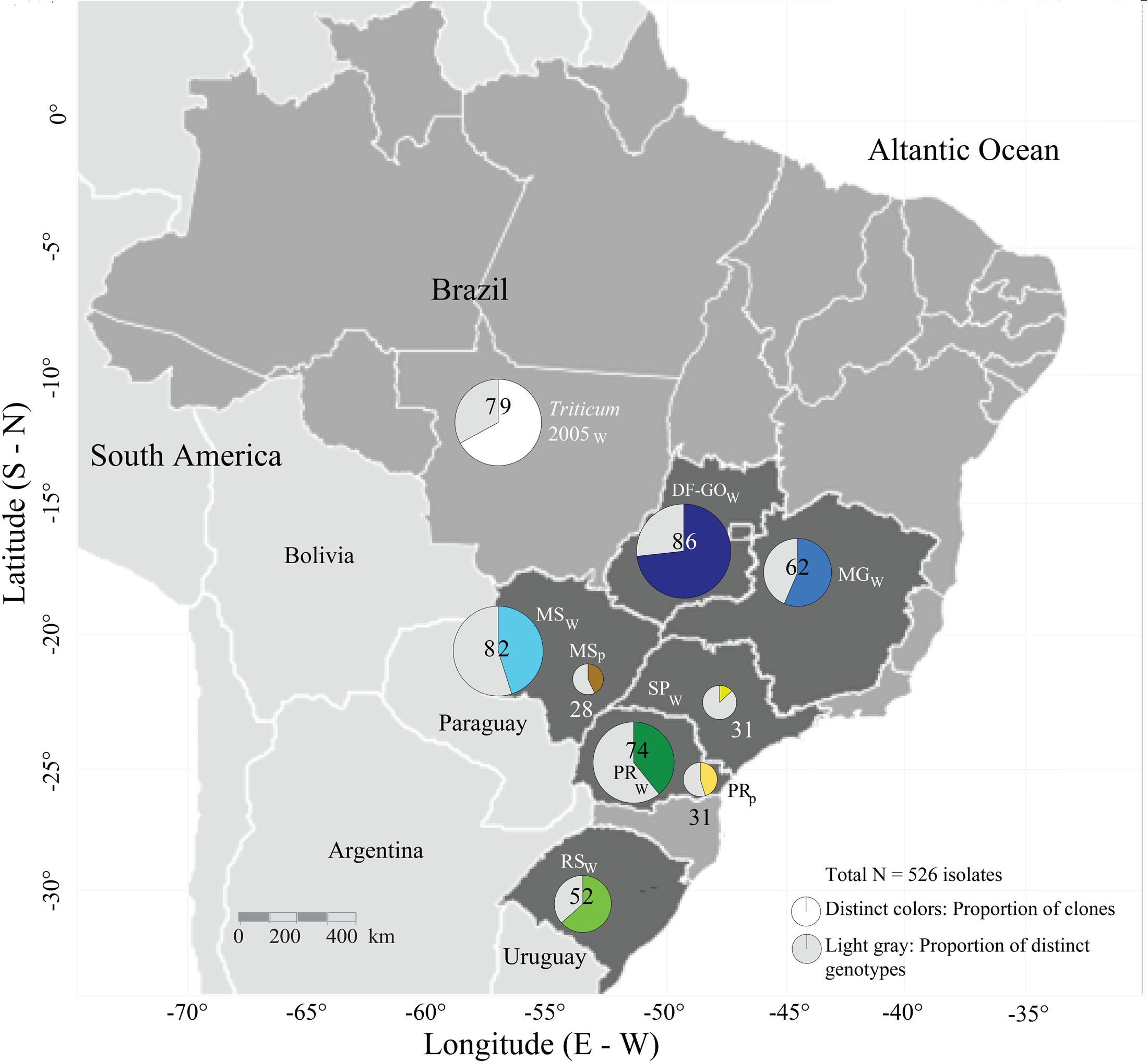
Geographical location of populations of *Pyricularia graminis-tritici* and *P. oryzae* examined in this study. The distinct colors in each population indicate the proportion of clones, while light gray indicates the proportion of distinct genotypes. Population 2005w was included because it represents a collection of MLMG genotypes sampled earlier in 2005 from central-southern Brazil.

**Table 3.** Number of multilocus microsatellite genotypes shared between Brazilian population of *Pyricularia graminis-tritici* from wheat and ther poaceous hosts.

| Species, populations | Wheat (*Triticum aestivum*) |  |  |  |  |  |  | Other Poaceae |  |
| --- | --- | --- | --- | --- | --- | --- | --- | --- | --- |
|  | 2005<sub>w</sub> | DF-GO<sub>w</sub> | MG<sub>w</sub> | MS<sub>w</sub> | PR<sub>w</sub> | RS<sub>w</sub> | SP<sub>w</sub> | MS<sub>p</sub> | PR<sub>p</sub> |
| 2005<sub>w</sub> | - | 0 | 0 | 0 | 0 | 0 | 0 | 0 | 0 |
| DF-GO<sub>w</sub> |  | - | 9 | 1 | 0 | 1 | 0 | 0 | 0 |
| MG<sub>w</sub> |  |  | - | 3 | 1 | 4 | 0 | 0 | 0 |
| MS<sub>w</sub> |  |  |  | - | 8 | 8 | 2 | 5 | 4 |
| PR<sub>w</sub> |  |  |  |  | - | 1 | 4 | 0 | 3 |
| RS<sub>w</sub> |  |  |  |  |  | - | 0 | 3 | 0 |
| SP<sub>w</sub> |  |  |  |  |  |  | - | 0 | 1 |
| MS<sub>p</sub> |  |  |  |  |  |  |  | - | 3 |
| PR<sub>p</sub> |  |  |  |  |  |  |  |  | - |
| Total number of shared genotypes in each population | 0 | 9 | 11 | 14 | 10 | 10 | 5 | 7 | 7 |
| Total number of isolates with shared genotypes in each population | 0 | 68 | 45 | 43 | 19 | 41 | 8 | 12 | 21 |

**Table 4.**
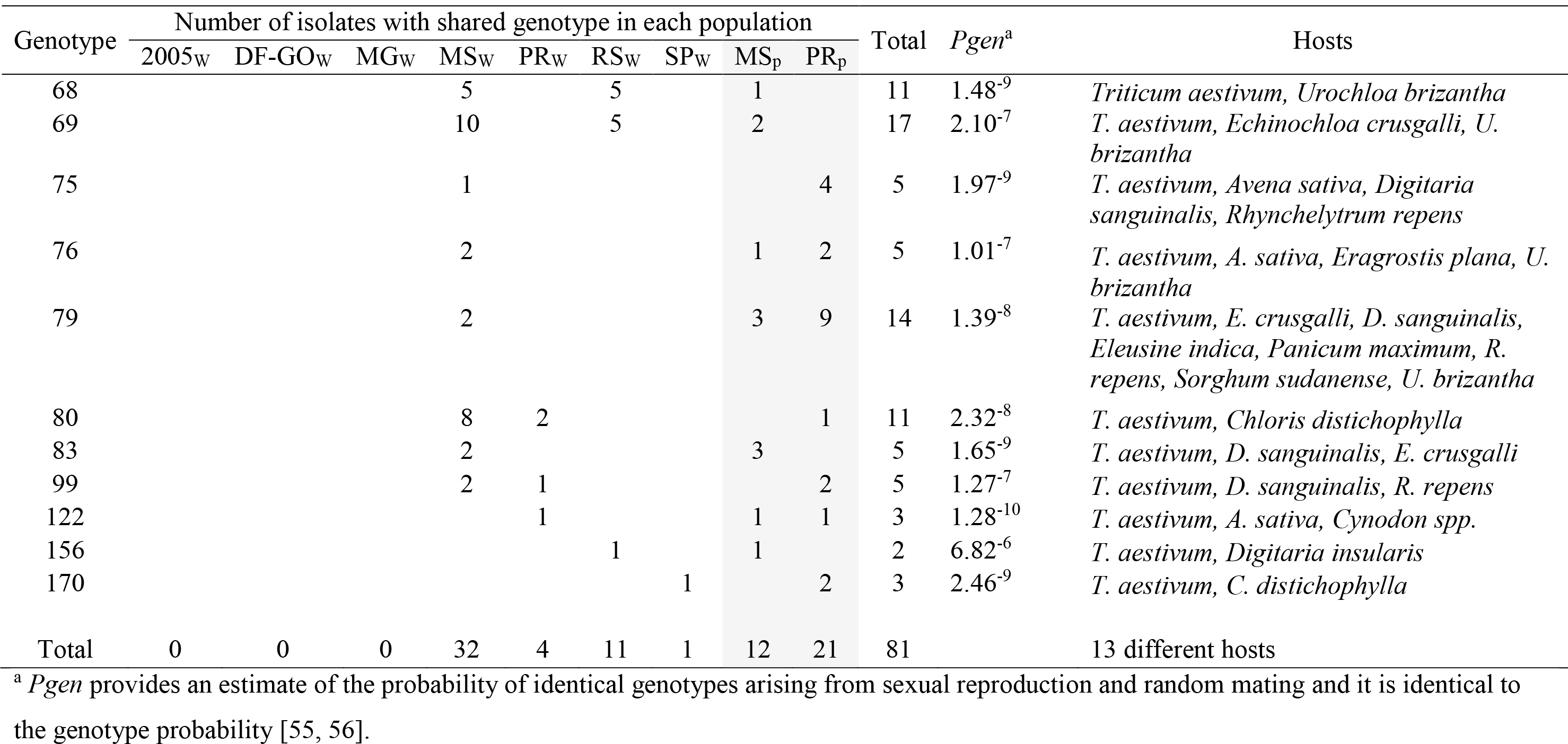
Number of isolates showing each of the eleven multilocus microsatellite genotypes shared among sympatric populations of *Pyricularia graminis-tritici* sampled from wheat and other poaceous hosts from Central-southern Brazil

The MS_P_ and PR_P_ populations sampled from other grass hosts shared 11 MLMGs with the populations from wheat. These 11 shared MLMGs were found in 48 strains originating from wheat and 33 strains recovered from other grass species, including *Avena sativa, Chloris distichophylla, Cynodon* spp., *Digitaria insularis, Digitaria sanguinalis, Echinochloa crusgalli, Eleusine indica, Eragrostis plana, Panicum maximum, Rhynchelytrum repens, Sorghum sudanense and Urochloa brizantha* (Table 4). The genetic similarity among all MLMGs and their geographical and host distributions are displayed as a minimal spanning network in Fig 3, with the 11 shared MLMGs indicated in red text. The probability that any two isolates drawn at random from the pool of 526 isolates would share one of these 11 MLMGs by chance in a recombining population ranged from 6.82^−6^ to 1.28^−10^ [55, 56] (Table 4), hence it is highly likely that isolates with the same MLMG represent the same clone or clonal lineage. These 11 MLMGs found on both wheat and other grasses provide compelling evidence for the existence of *Pygt* clones with a broad host range, with transmission among hosts growing in the same region likely occurring via dispersal of asexual spores, and transmission among distant geographical regions likely occurring via movement on infected seeds.

**Fig 3.**
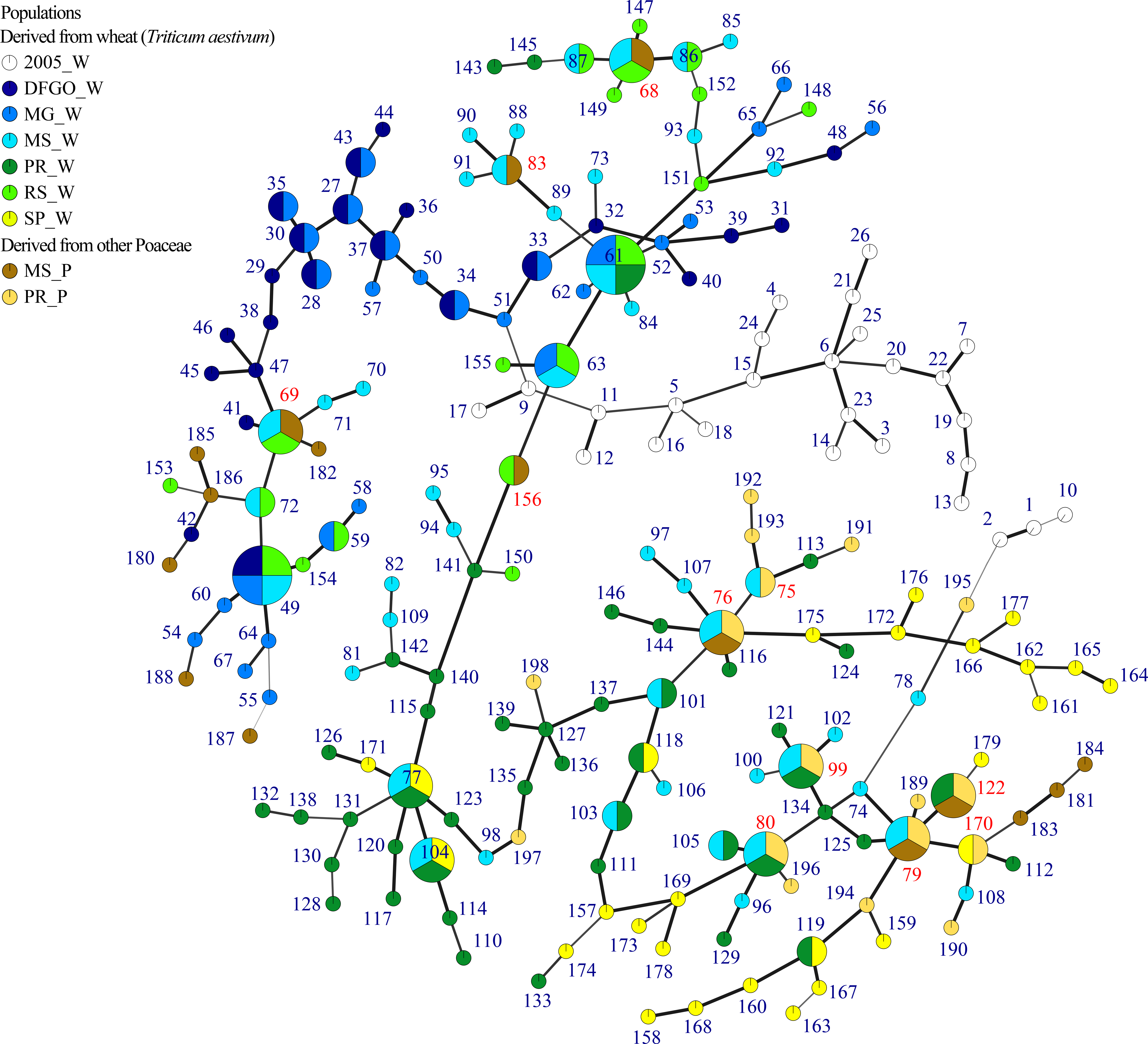
Minimum spanning network based on Bruvo distance for comparing 219 multilocus microsatellite genotypes (MLMG) of *Pyricularia graminis-tritici* isolates obtained from wheat and other poaceous hosts, and *P. oryzae* obtained from rice. Each node in the network represents a single haploid MLMG determined using 11 microsatellite loci. The size of the node (circle) represents the frequency of the sampled MLMGs. The shading (colors) of the nodes represents the membership of each population, while the thickness of the connecting lines and shading represent the degree of relationship between MLMGs. The line length is arbitrary. MLMGs shared among populations of *P. graminis-tritici* from wheat and other grasses are shown in red, while MLMGs associated only with one host are showed in black.

The clonal fraction inferred in each geographical population ranged from 0.13 in SP_w_ to 0.72 in DF-GO_w_, whereas the evenness ranged from 0.19 in the DF-GO_w_ population to ~ 0.90 in SP_w_. Overall, we found that the MLMGs were not uniformly distributed in the majority of the populations (Table 2). The effective number of genotypes (*G*_*o*_) ranged from 4.5 to 23.4 and was highest in *Pygt* populations from SP_w_ (*G*_*o*_ = 23.4), MS_w_ (*G*_*o*_ = 21.8) and PR_w_ (*G*_*o*_ = 18.3) and lowest in DF-GO_w_ (*G*_*o*_ = 4.5) (Table 2). The allelic richness averaged across ten populations was 2.75. The MS_P_ population from other grasses had the highest allelic richness (3.18) (Table 2).

### *Pygt* populations on wheat and other grasses are connected by gene flow

The overall fixation index indicated a weak but significant differentiation (*R*_*ST*_ = 0.07, *p* ≤ 0.001) among all populations. When *Pygt* populations from wheat were analyzed separately, AMOVA showed a low but still significant level of population differentiation (*R*_*ST*_ = 0.07, *p* ≤ 0.001), with 93% of the genetic variation distributed within populations. In contrast, when the two *Pygt* populations from other grasses (separated by ~ 430 km) were compared, AMOVA indicated an absence of population differentiation (*R*_*ST*_ = 0.02, *p* = 0.29), with 98% of genetic variation distributed within grass-infecting populations. The orthogonal contrast of *Pygt* populations from wheat with *Pygt* populations from other poaceous hosts was significant but the level of differentiation was very low (*R*_*CT*_ = 0.04), with the majority of genetic variation distributed within populations (93%) (Table 5). It is notable that no subdivision was found for 12 of the 15 pairwise comparisons between the two *Pygt* populations obtained from other grass hosts (MS_P_ and PR_P_) and the *Pygt* populations from wheat (Table 6).

**Table 5.**
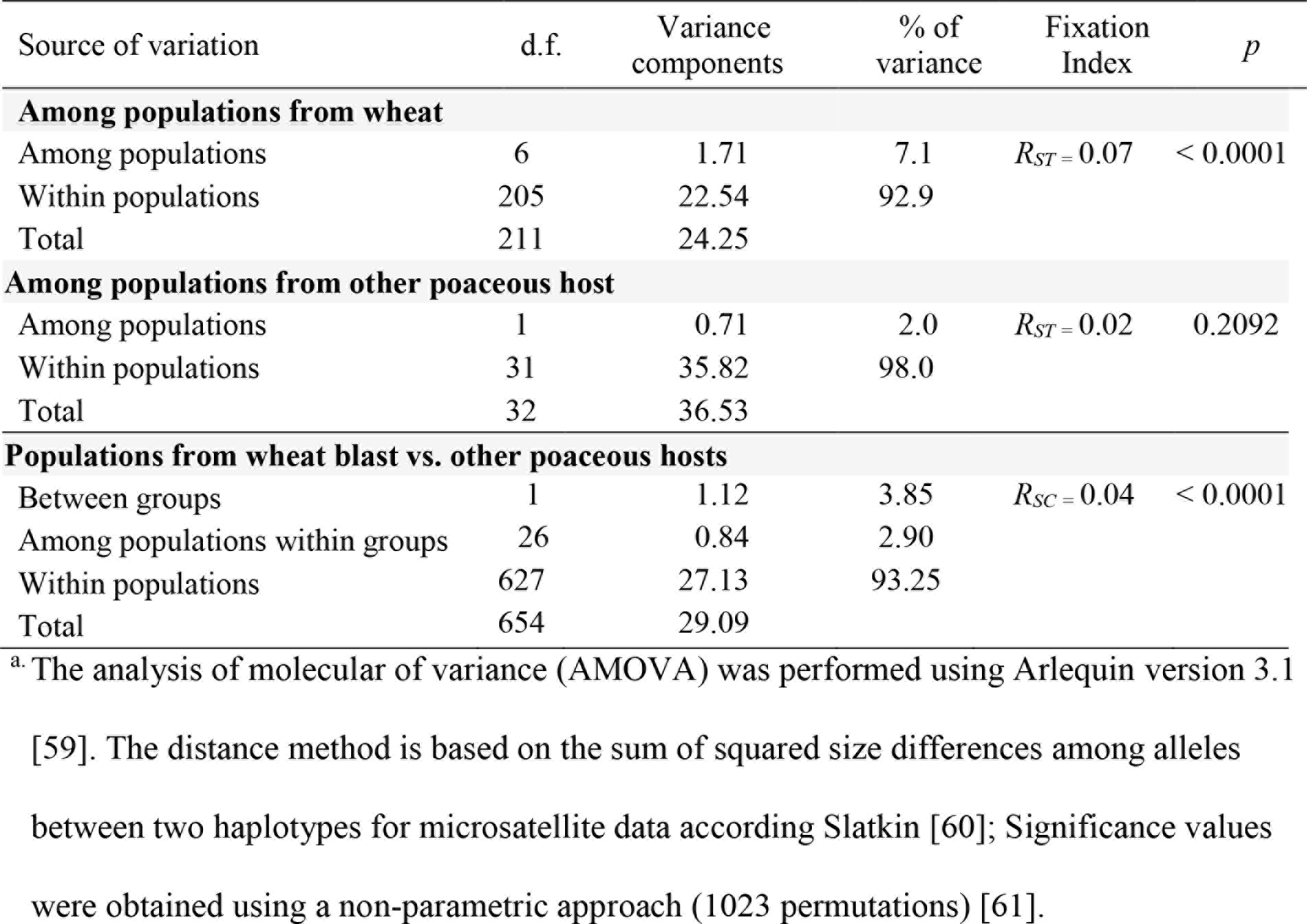
Hierarchical distribution of gene diversity among populations of *Pyricularia graminis-tritici* from wheat and other poaceous hosts and *P. oryzae* from rice in Central-southern Brazil ^a^.

**Table 6.** Pairwise differentiation among populations of *Pyricularia graminis-tritici* from wheat and other poaceous hosts in Central-southern Brazil

| $R_{ST}$ | Wheat (*T. aestivum*) | | | | | | | Other Poaceae | |
| --- | --- | --- | --- | --- | --- | --- | --- | --- | --- |
|  | 2005<sub>W</sub> | DF-GO<sub>W</sub> | MG<sub>W</sub> | MS<sub>W</sub> | PR<sub>W</sub> | RS<sub>W</sub> | SP<sub>W</sub> | MS<sub>P</sub> | PR<sub>P</sub> |
| 2005<sub>W</sub> | - |  |  |  |  |  |  |  |  |
| DF-GO<sub>W</sub> | 0.0066 | - |  |  |  |  |  |  |  |
| MG<sub>W</sub> | 0.0078 | 0.0340 | - |  |  |  |  |  |  |
| MS<sub>W</sub> | 0.0561 | 0.1567* | 0.0403 | - |  |  |  |  |  |
| PR<sub>W</sub> | 0.0944* | 0.2387* | 0.1213* | 0.0092 | - |  |  |  |  |
| RS<sub>W</sub> | 0.0610 | 0.1583* | 0.0400 | 0.0298 | 0.0830 | - |  |  |  |
| SP<sub>W</sub> | 0.0281 | 0.1734* | 0.0969 | 0.0480 | 0.0660 | 0.1499 | - |  |  |
| MS<sub>P</sub> | -0.0099 | 0.0750 | 0.0203 | 0.0316 | 0.0690 | 0.0733 | -0.0050 | - |  |
| PR<sub>P</sub> | 0.0380 | 0.1794* | 0.1274* | 0.0927 | 0.0674 | 0.1661* | 0.0044 | 0.0196 | - |

<sup>a</sup>. Fixation index among the nine populations of *P. graminis-tritici*: $R_{ST} = 0.07$ ( $p \leq 0.001$ )
<sup>b</sup>. Calculation were conducted on clone corrected data and based on the sum of squared size differences among alleles ( $R_{ST}$ ) between two
haplotypes for microsatellite data according Slatkin [60]. The test was performed using Arlequin version 3.1[59]. Significance values have
been tested using a non-parametric approach (1023 permutations) [61], at $\alpha = 0.05$ after Bonferroni correction for multiple comparisons [62].
$R_{ST}$ values followed by ‘\*’ showed statistically significant $p$ -values at corrected $\alpha = 0.005$ ).

### Historical gene flow was detected among *Pygt* populations from wheat and other grasses

The unidirectional migration models gave a better fit to the data than the panmictic or bidirectional models (Table 7). Historical migration analyses support unidirectional gene flow into the *Pygt* population infecting wheat from the *Pygt* population infecting other grasses (contributing 4.3 migrants per generation in average) (Table 8), suggesting that the *Pygt* population infecting wheat is composed of immigrants from the *Pygt* population infecting other grasses. There were no significant differences between Θ values (Table 8).

**Table 7.**
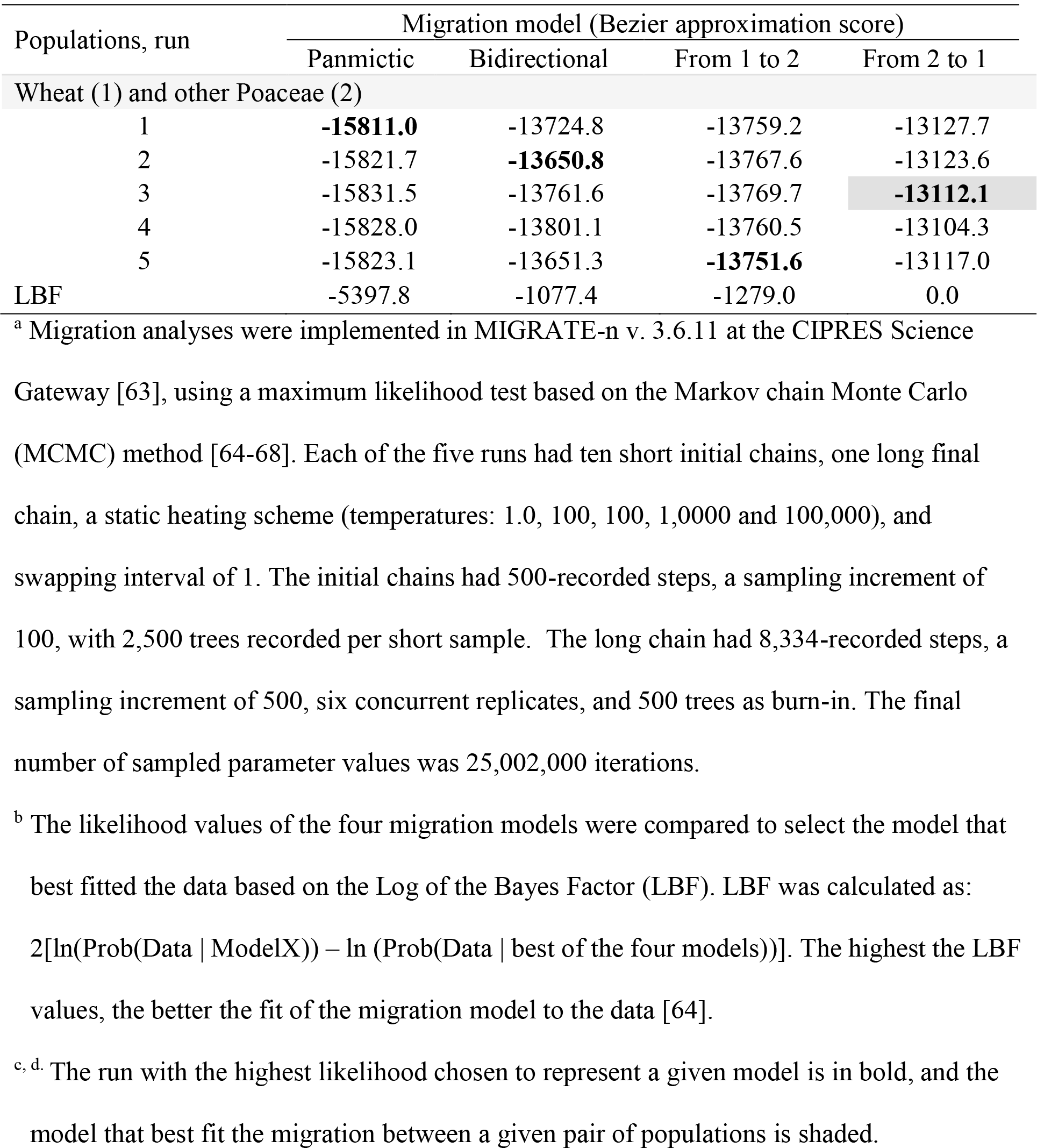
Comparison of models of historical migration between pairs of Brazilian populations of *Pyricularia graminis-tritici* grouped by original hosts (wheat and other Poaceae) based on Bezier approximation scores to the marginal likelihood^a^

**Table 8.** Migration parameter between pairs of Brazilian populations of *Pyricularia graminis-tritici* from wheat and other Poaceae, under the best fit migration model and obtained by Bayesian inference.

| Population pairs | Migration model that best fit the data | Migration parameters estimates <sup>a</sup> |  |  |
| --- | --- | --- | --- | --- |
| | | $\Theta_{\text{Wheat}}^b$ | $\Theta_{\text{Other Poaceae}}$ | $xNm_{\text{Donor} \rightarrow \text{Recipient}}^c$ |
| Wheat and other Poaceae | Directional (from other Poaceae to wheat) | 2.97<br>(0.0001 – 3.48) | 0.63<br>(0.0001 – 0.84) | $xNm_{\text{Poaceae} \rightarrow \text{Wheat}} = 4.26$<br>(0.0001 - 13.00) |

<sup>a</sup> Bayesian estimates of the migration parameters were calculated with MIGRATE-n v. 3.6.11 at the CIPRES Science Gateway [63],
using a maximum likelihood test based on the Markov chain Monte Carlo (MCMC) method [64-68]. Values represent the mean
Bayesian estimate, and are followed by the 95% credibility intervals for each parameter given by the 0.025 and 0.975 quantiles of its *a posteriori* distribution in parenthesis.
<sup>b</sup> Theta ( $\Theta$ ) values provide a measure of the effective population size; for haploids, $\Theta = 2Ne\mu$ , where $Ne$ = effective population size and $\mu$ = mutation rate for each locus [68].
<sup>c</sup> $xNm_{\text{Donor} \rightarrow \text{Recipient}}$ is the number of immigrants per generation; where $N$ is the real population size, $m$ is the fraction of the new
immigrants of the population per generation, and $x$ is an inheritance scalar and $x=1$ for haploids. The number of immigrants per
generation can also be expressed as the product $\Theta M$ ( $xNm_{\text{Donor} \rightarrow \text{Recipient}} = \Theta_{\text{recipient}} M_{\text{Donor} \rightarrow \text{Recipient}}$ ); where $M$ is $m$ divided by the mutation rate $\mu$ ( $M = m/\mu$ ), and it represents the importance of variability brought into the receiving population by immigration from the donor
population compared with the variability created by mutation [68].

### Most of the *Pygt* populations were sexually recombining

We consider a population to be sexually recombined when the majority of locus pairs are at gametic equilibrium and/or *I*_*A*_ or 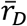 are not significant (*p* > 0.05). Under these assumptions, 7 of the 9 populations had signatures consistent with sexual recombination. Only 2005w and MSw showed evidence for significant clonal reproduction, with six and five pairs of loci showing significant GD, respectively, and significant *I*_*A*_ and 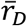 (*p* < 0.001) (Table 9).

**Table 9.** Estimates of gametic disequilibrium in populations of *Pyricularia graminis-tritici* from wheat and other poaceous hosts in Brazil.

| Population | Number of monomorphic loci | Locus pairs at significant disequilibrium<sup>a</sup> | Bonferroni correction $\alpha^b$ | *Mat1-1* (%) | *Mat1-2* (%) | Ratio *Mat1-1*:<br>*Mat1-2* | $I_A^c$ | $\bar{r}_D^c$ | $p^c$ |
| --- | --- | --- | --- | --- | --- | --- | --- | --- | --- |
| 2005<sub>w</sub> | 3 | 5 of 28 | 0.00183 | 81.0 | 19.0 | 4:1 | 1.818 | 0.263 | < **0.001** |
| DF-GO<sub>w</sub> | 1 | 0 of 45 | 0.00114 | 100.0 | 0.0 | 1:0 | -0.191 | -0.022 | 0.850 |
| MG<sub>w</sub> | 2 | 1 of 36 | 0.00142 | 100.0 | 0.0 | 1:0 | 0.177 | 0.022 | 0.097 |
| MS<sub>w</sub> | 1 | 6 of 45 | 0.00114 | 77.8 | 22.2 | 4:1 | 0.648 | 0.073 | < **0.001** |
| PR<sub>w</sub> | 1 | 0 of 45 | 0.00114 | 90.9 | 9.1 | 10:1 | -0.093 | -0.011 | 0.777 |
| RS<sub>w</sub> | 2 | 1 of 36 | 0.00142 | 73.7 | 26.3 | 3:1 | 0.041 | 0.005 | 0.365 |
| SP<sub>w</sub> | 3 | 0 of 28 | 0.00183 | 96.3 | 3.7 | 26:1 | 0.039 | 0.006 | 0.347 |
| MS<sub>p</sub> | 1 | 0 of 45 | 0.00114 | 23.1 | 76.9 | 1:4 | 0.694 | 0.079 | **0.001** |
| PR<sub>p</sub> | 3 | 0 of 28 | 0.00183 | 66.7 | 33.3 | 2:1 | -0.079 | -0.012 | 0.608 |

<sup>a</sup> Pairs of loci at significant disequilibrium according to the Fisher exact test (probability test) implemented by GENEPOP 3.4 [69] at $p \leq 0,05$ after Bonferroni correction for multiple comparisons [62].
<sup>b</sup> Value of $\alpha$ after Bonferroni correction used for multiple comparisons in the calculation of locus pairs at significant disequilibrium. Initial significance $\alpha = 0.05$ .
<sup>c</sup> $I_A$ and $\bar{r}_D$ are indexes of multilocus gametic disequilibrium (for the random association of alleles among distinct locus pairs). $\bar{r}_D$ is adjusted for the number of loci. The calculation of $I_A$ and $\bar{r}_D$ and their significance was performed using Multilocus software according Agapow and Burt, [70]. We tested $H_0$ = complete panmixia based on 1,000 randomizations; if $p \leq 0.05$ the population is under significant disequilibrium.

Because MSw possessed the highest number of shared MLMGS among populations (N=32, Table 4), we believe that the GD detected in this case was generated by the large influx of immigrants into this population.

### Perithecia of *Pygt* develop on senescing stems of wheat and other grasses

To better understand the role of sexual reproduction in the *Pygt* life cycle and determine whether the sexual cycle was more likely to occur on wheat or other grass hosts we performed a fruiting experiment and measured the production of proto-perithecia (the primordium that when fertilized develops into a perithecium) and perithecia on different host substrates. The ascocarps formed on autoclaved pieces of wheat stem were indistinguishable from those observed on naturally senescing pieces of stems of wheat and other Poaceae. The proto-perithecia and perithecia developed on the epidermal plant surface and within stems, where they were partially immersed in the internode culm. Proto-perithecia were black or very dark brown and sub-globose shaped. The mature perithecia were black and generally formed long beaks that often came from perithecia that were immersed in the plant tissue (Fig 4 and 5). Perithecia showed a mean size of 196 μm in length and 128 μm in width, with average neck size of 243 μm in length and 27 μm in width. Only the proto-perithecia formed on *Phalaris canariensis* reached a mature size consistent with complete development (Fig 5), suggesting that the sexual cycle was more likely to be completed on *P. canariensis* than on wheat.

**Fig 4.**
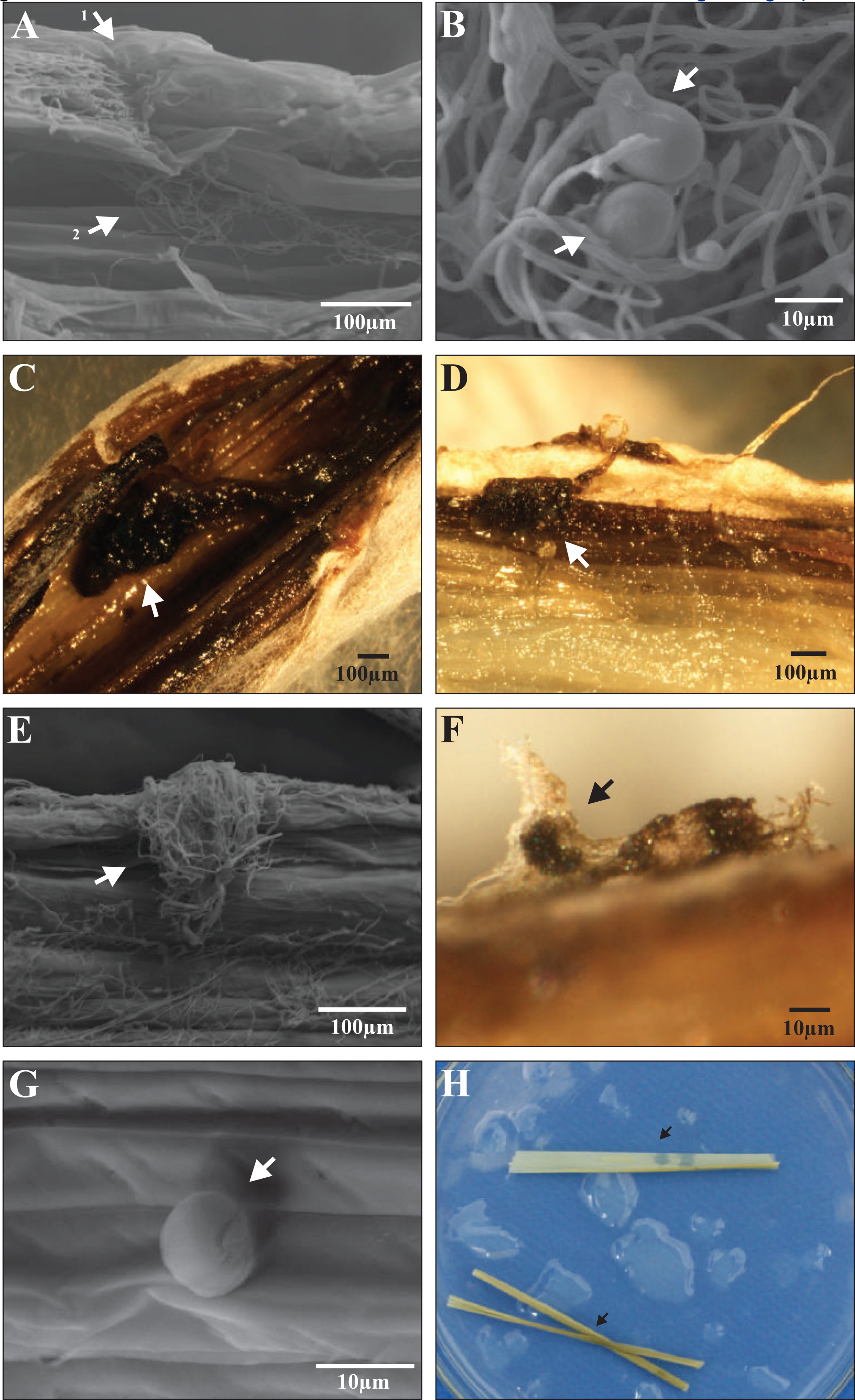
Development of proto-perithecia and perithecia of *Pyricularia graminis-tritici* induced by injection of living conidia of isolates PY33.1 (*Mat1-1*) and PY05046 (*Mat1-2*) within autoclaved stems sections of wheat (*Triticum aestivum*) cv. MGS Brilhante. Panel A, site of injection (arrow 1) and fungal colonization within plant tissues (arrow 2); development of proto-perithecia (B) and perithecia (C) inside stems; D, perithecia developing from the internal plant tissues to beak emersion; proto-perithecia at interface (E) and on surface of plant tissues (F and G). H, Control composed of autoclaved stems without inoculation. The images of panels A, B, E and G were acquired by scanning electron microscope. Images of panels C, D and F were acquired by light microscope.

**Fig 5.**
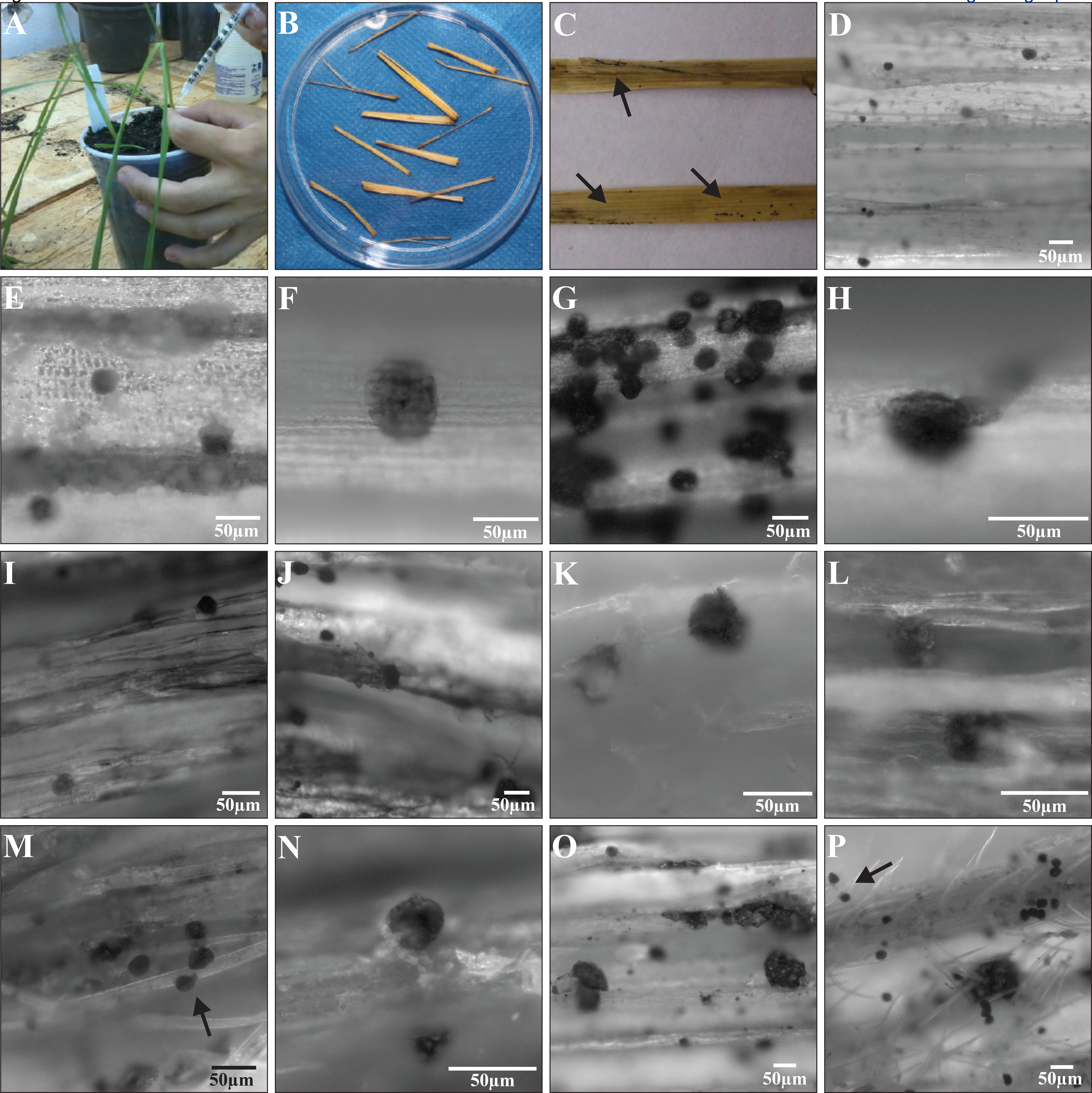
Development of proto-perithecia and perithecia of *Pyricularia graminis-tritici* induced by injection of living conidia of isolates Py33.1 (*Mat1-1*) and Py5046 (*Mat1-2*) within senescing stems sections of different Poaceae species. Panel A, Procedure for inoculation by injection of living spores into the host stems. B, Pieces of stem placed at 120 mm Petri dishes to incubation in humid chamber 1 month after inoculation. C, Stems with proto-perithecia and/or perithecia development after incubation in humid chamber (arrows). Fruiting body in different plant species: D, canary seeds (*Phalaris canariensis*); E, rice (*O. sativa*) cv. Primavera; F, rice cv. Relampago; G, red rice (*O. sativa*) cv. Yin Lu 30; H, *Brachiaria* cv. Hybrid Mulato; I, barley (*Hordeum vulgare*) cv. BR Elis; J, barley cv. MN 743; K,Rye (*Secale cereale*) cv. BR1; L, black oat (*Avena strigosa*) cv. Embrapa 29 Garoa; M, foxtail millet (*Setaria italica*); N, wheat (*Triticum aestivum*) cv. BRS 264; O, wheat cv. MGS Brilhante; P, triticale (x *Triticosecale*) cv. IAC Caninde. The images of panels D to P were acquired by bright field microscopy.

### The virulence spectra of *Pygt* populations varied across geographical regions

We examined the virulence spectra for 173 *Pygt* isolates on both seedlings and detached heads of ten differential wheat cultivars and one barley cultivar. These differentials were chosen based on previous experiments which suggested a gene-for-gene interaction that would allow us to distinguish *Pygt* pathotypes [39]. Our aim in this analysis was to assess the geographical distribution of virulence groups of *Pygt* and determine if virulence groups were shared between strains infecting wheat and other grasses. The 173 assessed *Pygt* isolates, encompassing 80 unique MLMGs, produced typical leaf blast lesions (Fig. 6) and could be grouped into 25 seedling virulence groups (SVGs) (Table 10). These SVGs were named A to Y. SVG L was the predominant group, comprising 47% of the tested isolates. SVG A was the second most frequent group, found in 13% of tested isolates. The 23 remaining SVGs were relatively infrequent (Tables 10 and 11, Fig 7). SVG L was the most widely distributed virulence group across Brazil. The MSp population had the highest number of SVGs (11 groups), whereas the PRw and SPw populations had the lowest number of SVGs (1 and 2 groups, respectively). Nine SVGs (A, F to I, and K to N) were shared among *Pygt* isolates originating from wheat and other grasses (Tables 10 and 11).

**Fig 6.**
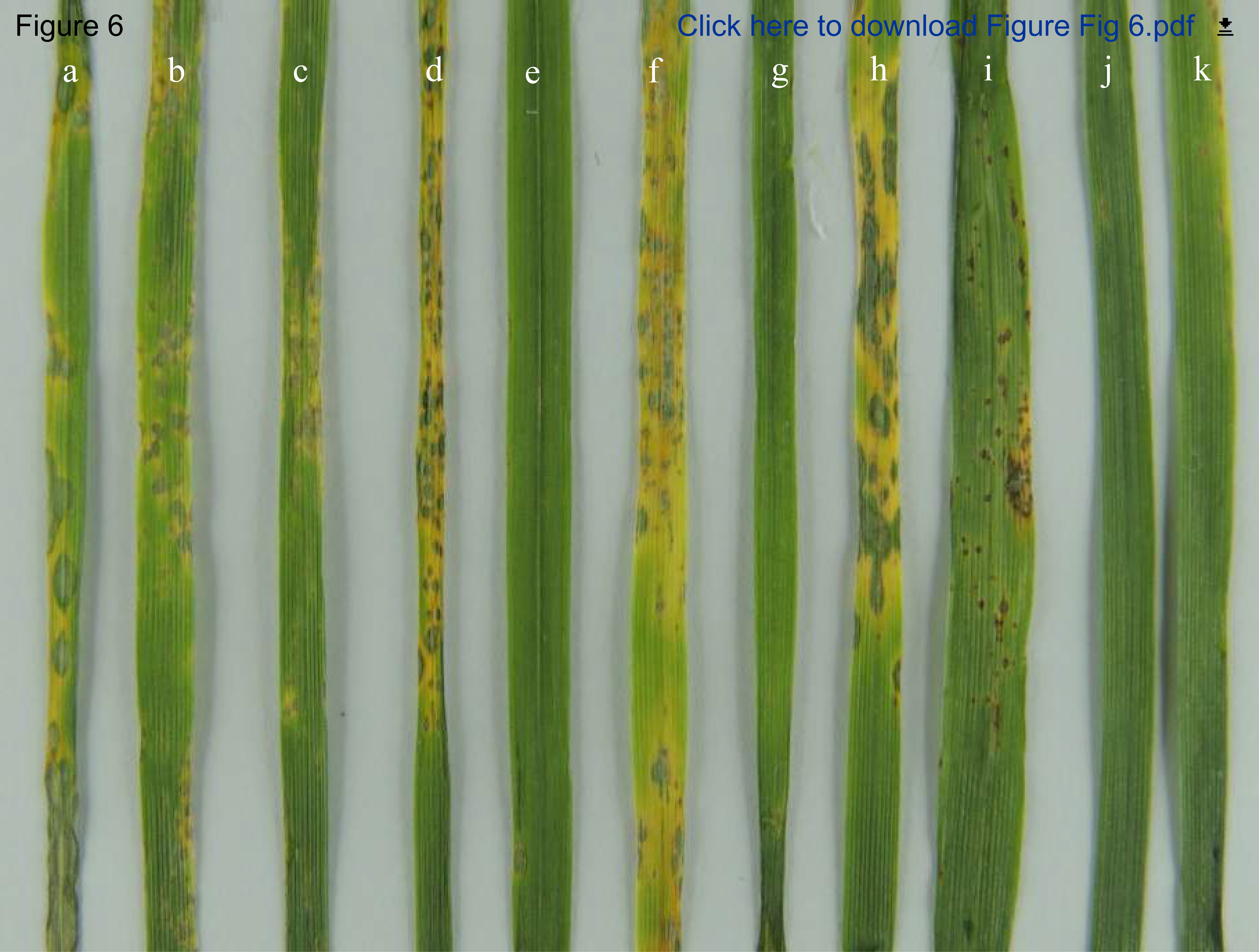
Virulence spectrum and typical blast lesions on wheat seedlings caused by isolates of *Pyricularia graminis-tritici* belonging to the predominant seedling virulence group (SVG L) on the differential set of ten wheat (*Triticum aestivum*) cultivars and one barley (*Hordeum vulgare*) cultivar. The differential set was consist of ten wheat cultivars: a) Anahuac 75; b) BR 18; c) BR 24; d) BRS 220; e) **BRS 229**; f) MGS 3 Brilhante; g), **BRS Buruti**; h) CNT 8; j) **Renan**; k) **BRS 234**; and one barley cultivar: i) PFC 2010123. Varieties indicated in bold showed resistant reaction. Isolate inoculated: 12.1.109.

**Fig 7.**
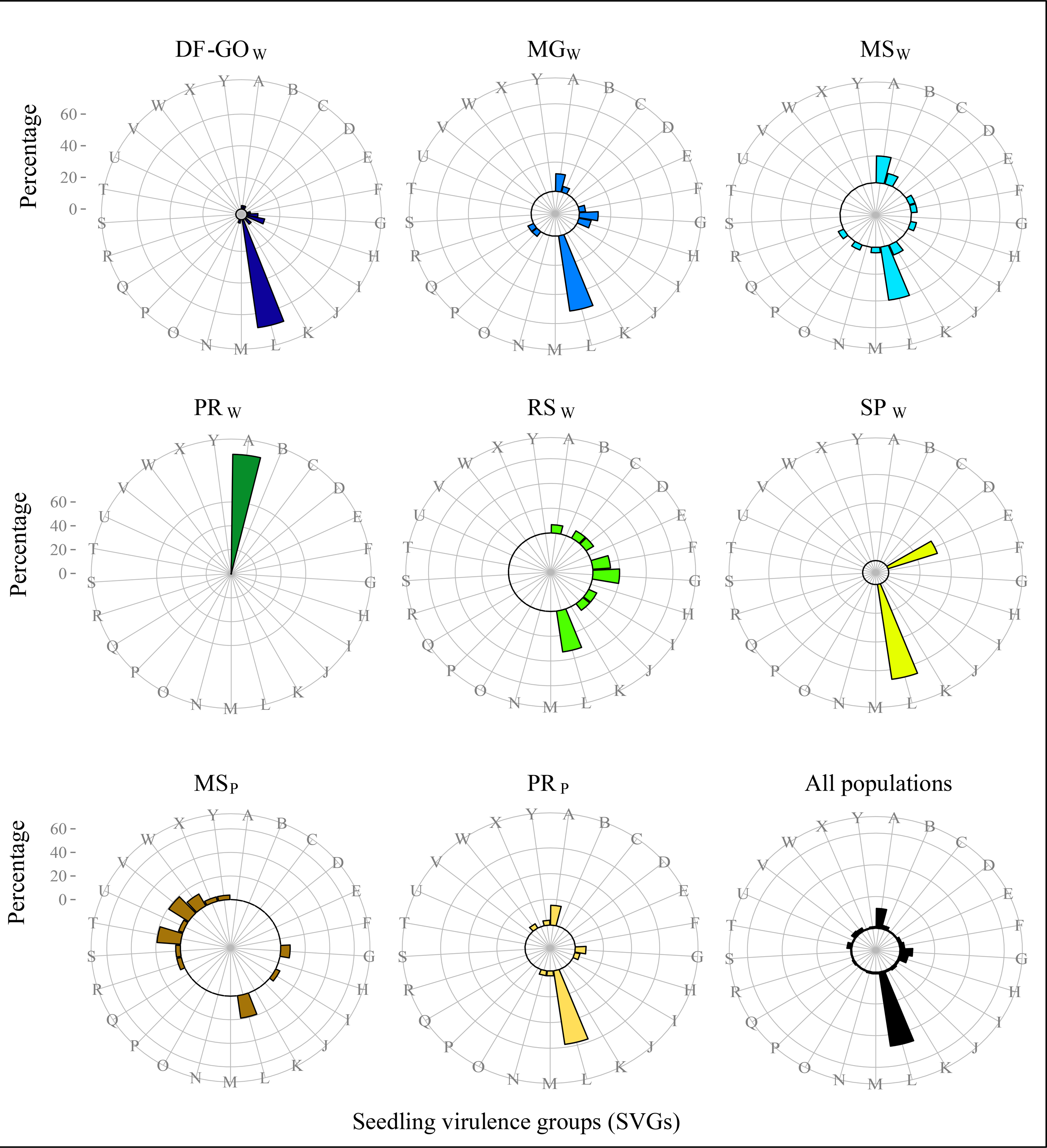
Distribution of seeding virulence groups (SVGs) of the wheat blast pathogen *Pyricularia graminis-tritici* in ten populations from central-southern Brazil.

**Table 10.**
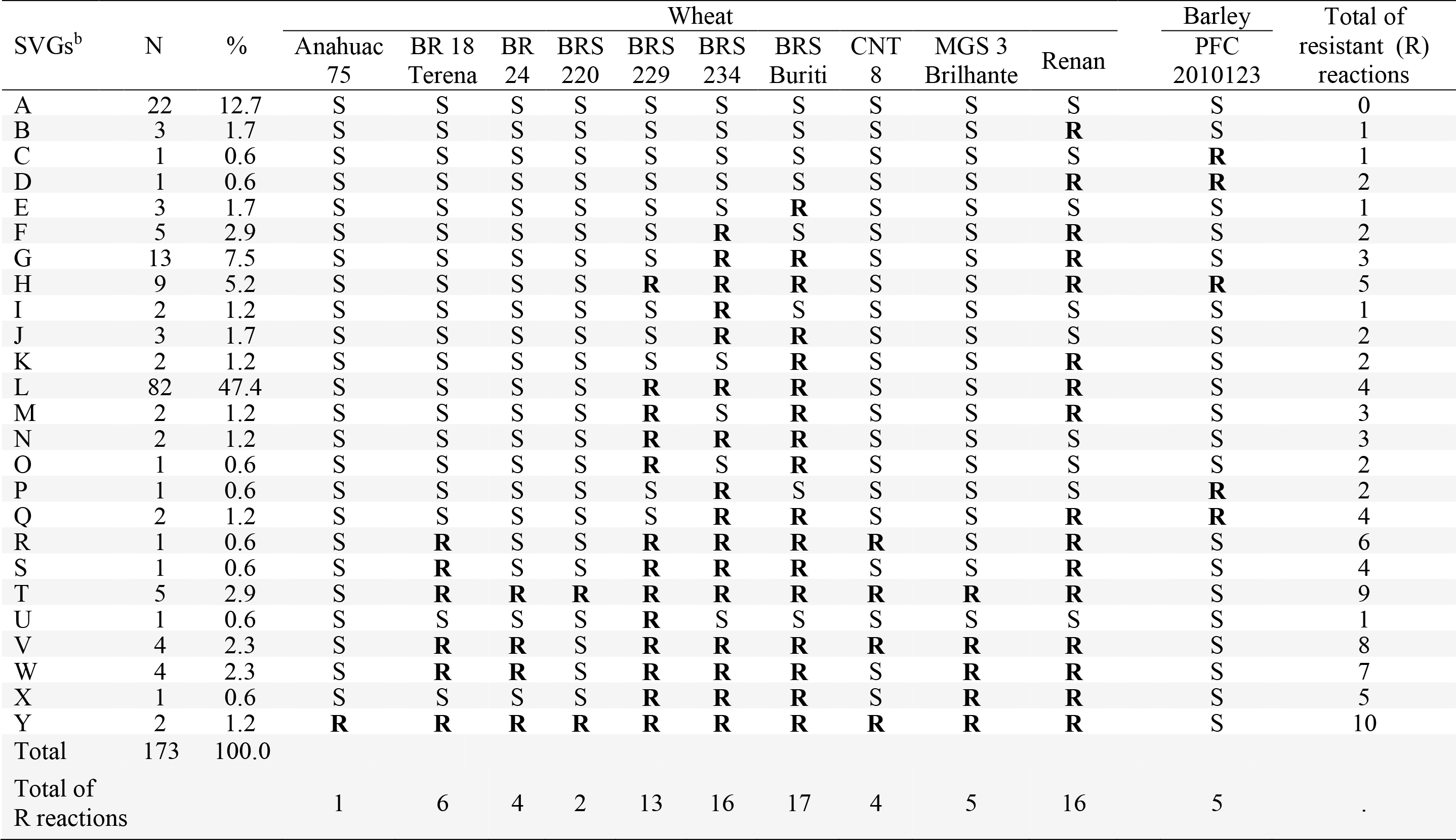
Pathogenicity spectra of seedling virulence groups (SVGs) of isolates of *Pyricularia graminis-tritici*^a^

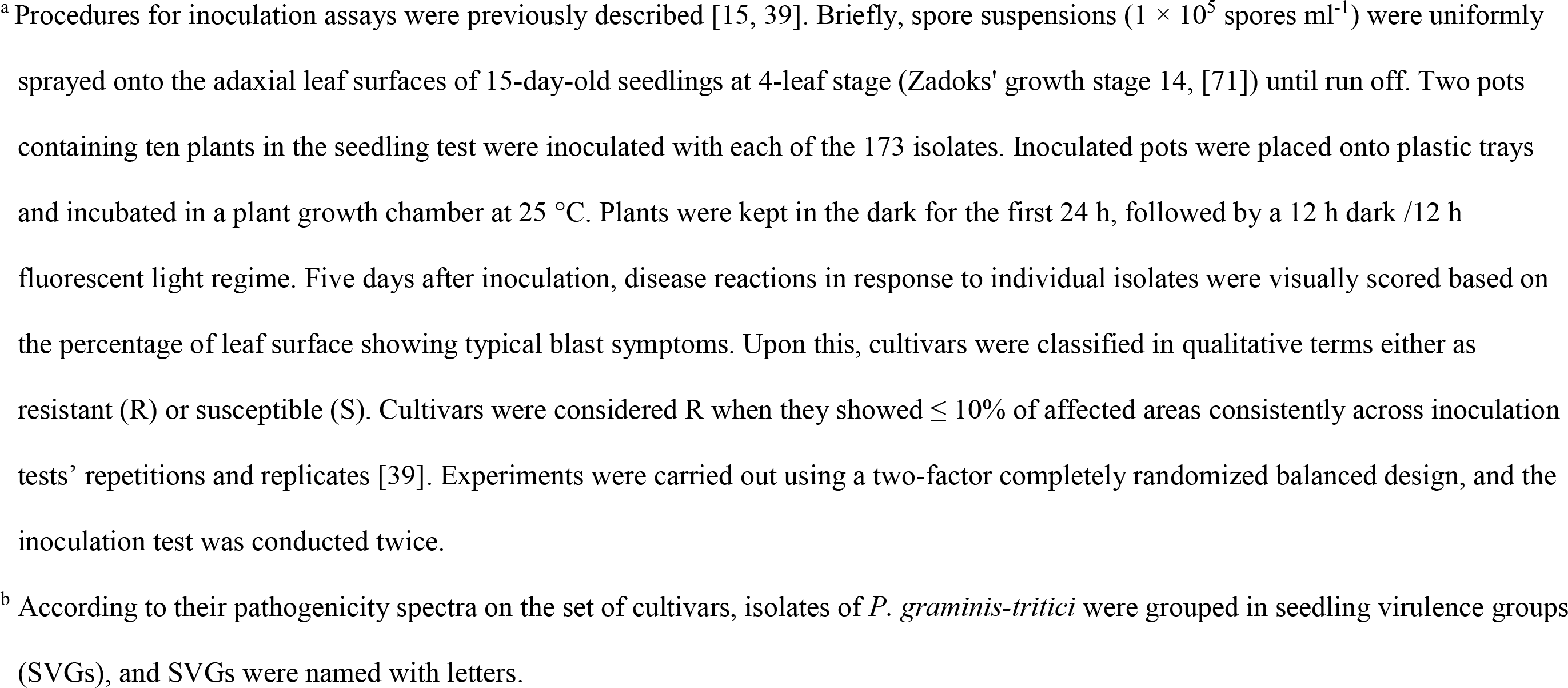

**Table 11.** Isolates of *Pyricularia graminis-tritici* assigned to each seedling virulence group (SVG) per population

| SVG | Population |  |  |  |  |  |  |  |  |  |  |  |  |  |  |  | Total |  |
| --- | --- | --- | --- | --- | --- | --- | --- | --- | --- | --- | --- | --- | --- | --- | --- | --- | --- | --- |
|  | DF_GO<sub>w</sub> |  | MG<sub>w</sub> |  | MS<sub>w</sub> |  | PR<sub>w</sub> |  | RS<sub>w</sub> |  | SP<sub>w</sub> |  | MS<sub>p</sub> |  | PR<sub>p</sub> |  |  |  |
|  | N | % | N | % | N | % | N | % | N | % | N | % | N | % | N | % | N | % |
| A | 1 | 2.3 | 3 | 12.5 | 5 | 20.0 | 8 | 100.0 | 1 | 6.7 | . | . | . | . | 4 | 15.4 | 22 | 12.7 |
| B | . | . | 1 | 4.2 | 2 | 8.0 | . | . | . | . | . | . | . | . | . | . | 3 | 1.7 |
| C | . | . | . | . | . | . | . | . | 1 | 6.7 | . | . | . | . | . | . | 1 | 0.6 |
| D | . | . | . | . | . | . | . | . | 1 | 6.7 | . | . | . | . | . | . | 1 | 0.6 |
| E | . | . | . | . | 1 | 4.0 | . | . | . | . | 2 | 33.3 | . | . | . | . | 3 | 1.7 |
| F | 1 | 2.3 | 1 | 4.2 | 1 | 4.0 | . | . | 2 | 13.3 | . | . | . | . | . | . | 5 | 2.9 |
| G | 3 | 6.8 | 3 | 12.5 | . | . | . | . | 3 | 20.0 | . | . | 2 | 8.0 | 2 | 7.7 | 13 | 7.5 |
| H | 5 | 11.4 | 2 | 8.3 | 1 | 4.0 | . | . | . | . | . | . | . | . | 1 | 3.8 | 9 | 5.2 |
| I | . | . | . | . | . | . | . | . | 1 | 6.7 | . | . | 1 | 4.0 | . | . | 2 | 1.2 |
| J | 2 | 4.5 | . | . | . | . | . | . | 1 | 6.7 | . | . | . | . | . | . | 3 | 1.7 |
| K | . | . | . | . | 2 | 8.0 | . | . | . | . | . | . | . | . | . | . | 2 | 1.2 |
| L | 31 | 70.5 | 12 | 50.0 | 10 | 40.0 | . | . | 5 | 33.3 | 4 | 66.7 | 5 | 20.0 | 15 | 57.7 | 82 | 47.4 |
| M | . | . | . | . | 1 | 4.0 | . | . | . | . | . | . | . | . | 1 | 3.8 | 2 | 1.2 |
| N | 1 | 2.3 | . | . | . | . | . | . | . | . | . | . | . | . | 1 | 3.8 | 2 | 1.2 |
| O | . | . | . | . | 1 | 4.0 | . | . | . | . | . | . | . | . | . | . | 1 | 0.6 |
| P | . | . | 1 | 4.2 | . | . | . | . | . | . | . | . | . | . | . | . | 1 | 0.6 |
| Q | . | . | 1 | 4.2 | 1 | 4.0 | . | . | . | . | . | . | . | . | . | . | 2 | 1.2 |
| R | . | . | . | . | . | . | . | . | . | . | . | . | 1 | 4.0 | . | . | 1 | 0.6 |
| S | . | . | . | . | . | . | . | . | . | . | . | . | 1 | 4.0 | . | . | 1 | 0.6 |
| T | . | . | . | . | . | . | . | . | . | . | . | . | 5 | 20.0 | . | . | 5 | 2.9 |
| U | . | . | . | . | . | . | . | . | . | . | . | . | 1 | 4.0 | . | . | 1 | 0.6 |
| V | . | . | . | . | . | . | . | . | . | . | . | . | 4 | 16.0 | . | . | 4 | 2.3 |
| W | . | . | . | . | . | . | . | . | . | . | . | . | 3 | 12.0 | 1 | 3.8 | 4 | 2.3 |
| X | . | . | . | . | . | . | . | . | . | . | . | . | 1 | 4.0 | . | . | 1 | 0.6 |
| Y | . | . | . | . | . | . | . | . | . | . | . | . | 1 | 4.0 | 1 | 3.8 | 2 | 1.2 |
| Total | 44 | 100.0 | 24 | 100.0 | 25 | 100.0 | 8 | 100.0 | 15 | 100.0 | 6 | 100.0 | 25 | 100.0 | 26 | 100.0 | 173 | 100.0 |
| Total SVGs | 7 | . | 8 | . | 10 | . | 1 | . | 8 | . | 2 | . | 11 | . | 8 | . | 25 | . |

The same isolates fell into nine different head virulence groups (HVGs) when virulence spectra were assessed on detached, mature wheat heads. Five of these HVGs (A to D, and T) had virulence spectra that were identical to the five SVGs (A to D, and T), so we used the same nomenclature for these SVGs and HVGs. The remaining HVGs were designated AA to DD. HVG A was the predominant virulence group, found in 138 isolates, followed by HVG B found in 25 isolates (Table 12). Both of these virulence groups were found in all *Pygt* populations (Table 13), including the grass-infecting populations. The remaining seven virulence groups were found in only 1 or 2 isolates. As found for the seedling assay, MSP was the population with the highest number of HVGs (6), and PRW was the population with the lowest number of HVGs (1) (Table 14, Fig 8 and 9).

**Table 12.** Pathogenicity spectra of head virulence groups (HVGs) of isolates of *P. graminis-triticf*^a^

| HVGs | N | % | Wheat |  |  |  |  |  |  |  |  |  | Barley | Total of<br>resistant (R)<br>reactions |
| --- | --- | --- | --- | --- | --- | --- | --- | --- | --- | --- | --- | --- | --- | --- |
|  |  |  | Anahuac<br>75 | BR 18<br>Terena | BR<br>24 | BRS<br>220 | BRS<br>229 | BRS<br>234 | BRS<br>Buriti | CNT<br>8 | MGS 3<br>Brilhante | Renan | PFC<br>2010123 |  |
| A | 138 | 79.8 | S | S | S | S | S | S | S | S | S | S | S | 0 |
| B | 25 | 14.5 | S | S | S | S | S | S | S | S | S | **R** | S | 1 |
| C | 2 | 1.2 | S | S | S | S | S | S | S | S | S | S | **R** | 1 |
| D | 2 | 1.2 | S | S | S | S | S | S | S | S | S | **R** | **R** | 2 |
| T | 1 | 0.6 | S | **R** | **R** | **R** | **R** | **R** | **R** | **R** | **R** | **R** | S | 9 |
| AA | 2 | 1.2 | S | S | **R** | S | S | S | S | S | S | S | S | 1 |
| BB | 1 | 0.6 | S | S | **R** | S | S | S | **R** | S | S | **R** | S | 3 |
| CC | 1 | 0.6 | S | S | S | S | S | **R** | S | S | S | **R** | **R** | 2 |
| DD | 1 | 0.6 | S | **R** | S | **R** | **R** | **R** | **R** | S | **R** | **R** | S | 7 |
| Total | 173 | 100.0 |  |  |  |  |  |  |  |  |  |  |  |  |
| Total of R reactions |  |  | 0 | 2 | 3 | 2 | 2 | 3 | 3 | 1 | 2 | 6 | 3 | - |

<sup>a</sup>. Procedures for inoculation assays have been previously described [15, 39]. In short, spore suspensions ( $1 \times 10^5$ spores ml<sup>-1</sup>) were uniformly sprayed onto the head detached heads harvested from plants between anthesis and initial grain-3 to grain-filling stage (Zadoks' growth stages 63 to 71, [71]) until run off. Three polyurethane foam blocks with ten detached heads apice were inoculated with each of the 173 isolates. Inoculated heads were placed in plastic boxes and incubated in a plant growth chamber at 25 °C. Plants were kept in the dark for the first 24 h, followed by a 12 h dark /12 h fluorescent light regime. Five days after inoculation, disease reactions in response to individual isolates were visually scored based on the percentage of detached head showing typical blast symptoms. Upon this, cultivars were classified in qualitative terms either as resistant (R) or susceptible (S). Cultivars were considered R when they showed $\leq 10\%$ of affected areas consistently across inoculation tests' repetitions and replicates [39]. Experiments were carried out using a two-factor completely randomized balanced design, and the inoculation test was conducted twice.
<sup>b</sup>. According to their pathogenicity spectra on the set cultivars, isolates of *P. graminis-tritici* were grouped in head virulence groups (HVGs). Groups were named with letters, and the name was maintained when the pathogenicity spectra on seedling and on detached heads coincided.

**Table 13.** Isolates of *Pyricularia graminis-tritici* assigned to each head virulence group (HVG) per population

| HVG | Population |  |  |  |  |  |  |  |  |  |  |  |  |  |  |  | Total |  |
| --- | --- | --- | --- | --- | --- | --- | --- | --- | --- | --- | --- | --- | --- | --- | --- | --- | --- | --- |
|  | DF_GO<sub>w</sub> |  | MG<sub>w</sub> |  | MS<sub>w</sub> |  | PR<sub>w</sub> |  | RS<sub>w</sub> |  | SP<sub>w</sub> |  | MS<sub>P</sub> |  | PR<sub>P</sub> |  |  |  |
|  | N | % | N | % | N | % | N | % | N | % | N | % | N | % | N | % | N | % |
| A | 33 | 75.0 | 17 | 70.8 | 23 | 92 | 8 | 100 | 10 | 66.6 | 5 | 83.3 | 19 | 76.0 | 23 | 88.3 | 138 | 79.8 |
| B | 10 | 22.7 | 6 | 25.0 | 2 | 8 | . | . | 3 | 20 | 1 | 16.7 | 2 | 8.0 | 1 | 3.9 | 25 | 14.5 |
| C | . | . | . | . | . | . | . | . | 1 | 6.7 | . | . | . | . | . | . | 2 | 1.2 |
| D | 1 | 2.3 | 1 | 4.2 | . | . | . | . | 1 | 6.7 | . | . | . | . | . | . | 2 | 1.2 |
| T | . | . | . | . | . | . | . | . | . | . | . | . | . | . | 1 | 3.9 | 1 | 0.6 |
| AA | . | . | . | . | . | . | . | . | . | . | . | . | 1 | 4.0 | 1 | 3.9 | 2 | 1.2 |
| BB | . | . | . | . | . | . | . | . | . | . | . | . | 1 | 4.0 | . | . | 1 | 0.6 |
| CC | . | . | . | . | . | . | . | . | . | . | . | . | 1 | 4.0 | . | . | 1 | 0.6 |
| DD | . | . | . | . | . | . | . | . | . | . | . | . | 1 | 4.0 | . | . | 1 | 0.6 |
| Total | 44 | 100.0 | 24 | 100.0 | 25 | 100.0 | 8 | 100.0 | 15 | 100.0 | 6 | 100.0 | 25 | 100.0 | 26 | 100.0 | 173 | 100.0 |
| Total HVGs | 3 | . | 3 | . | 2 | . | 1 | . | 4 | . | 2 | . | 6 | . | 4 | . | 9 | . |

**Fig 8.**
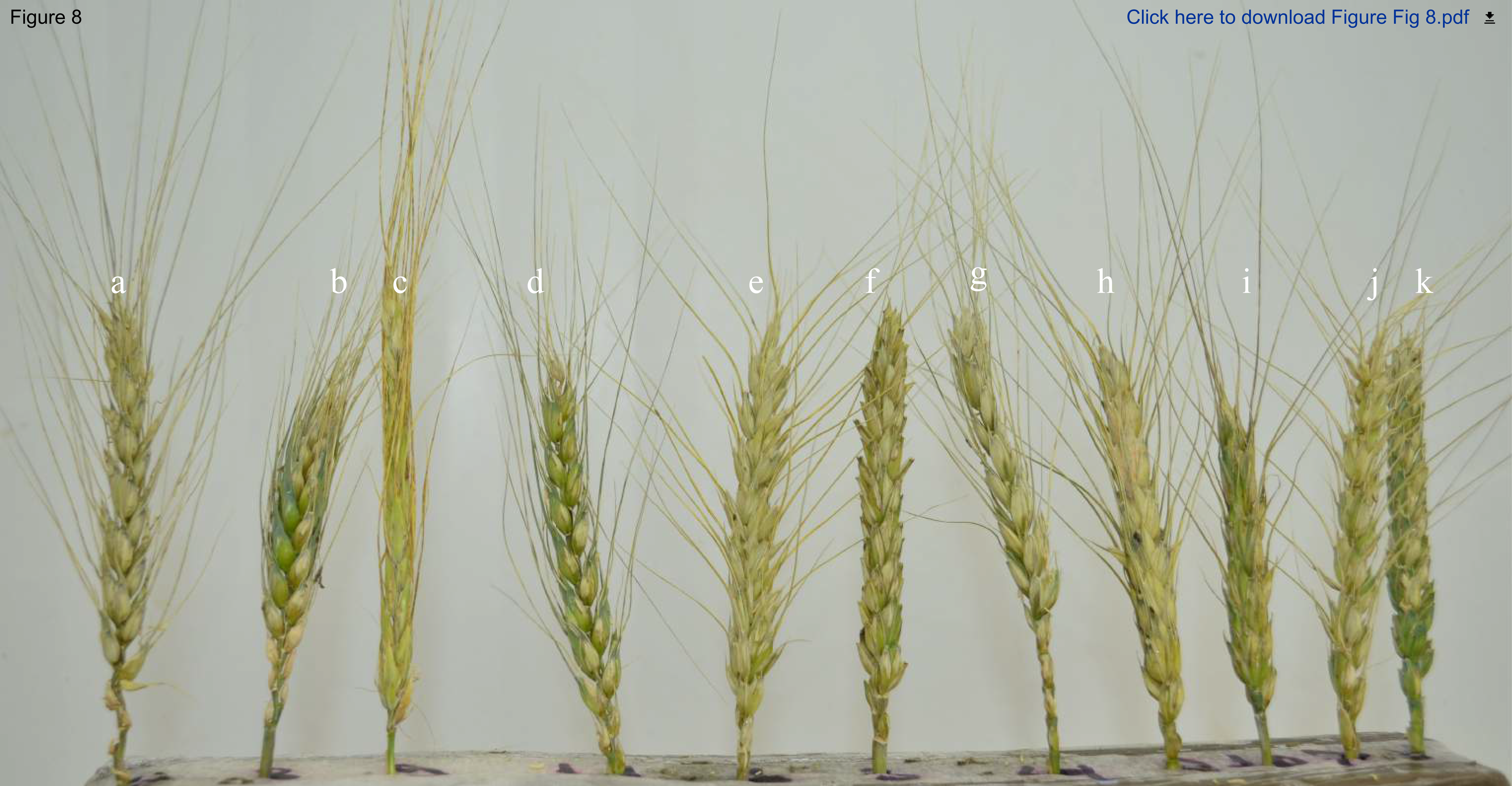
Virulence spectrum and typical blast lesions on wheat heads caused by isolates of *Pyricularia graminis-tritici* belonging to the predominant head virulence group (HVG A) on the differential set of ten wheat (*Triticum aestivum*) cultivars and one barley (*Hordeum vulgare*) cultivar. The differential set was consist of ten wheat cultivars: a) BRS 229; b) CNT 8; d) BR 234; e) Anahuac 75; f) BR 24; g) BRS 220; h), BR 18; i) Renan; j) BRS Buriti; k) MGS 3 Brilhante; and one barley cultivar: c) PFC 2010123. All cultivars showed susceptible reactions. Isolate inoculated: 12.1.170.

**Fig 9.**
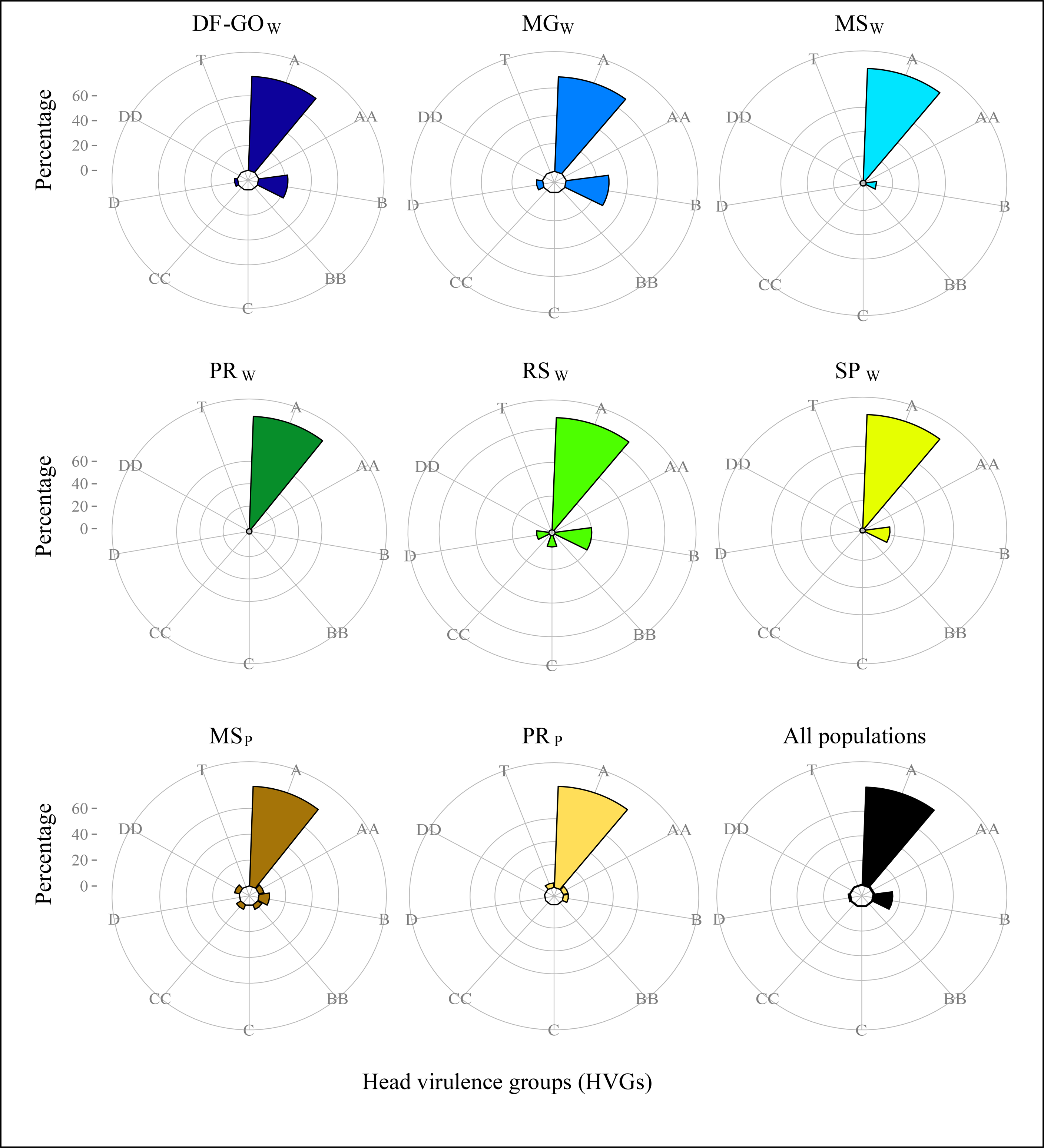
Distribution of head virulence groups (HVGs) of the wheat blast pathogen *Pyricularia graminis-tritici* in ten populations from central-southern Brazil.

## Discussion

The phylogenetic analyses based on entire genome sequences did not support the earlier hypothesis that two distinct species (named *P. graminis-tritici* (*Pygt*) and *P. oryzae* pathotype *Triticum* (*PoT*) in Fig 1) cause wheat blast [15]. Instead, our phylogenetic analyses indicate that *Pygt* is a single, highly diverse pathogen species with a broad host range that encompasses many grasses that were either native (e.g. *Chloris distichophylla, Cynodon* spp., *Digitaria insularis*) or introduced into Brazil for food production during the last 200 years. Our current phylogenetic analyses do not allow us to determine whether there were multiple origins or a single origin for the wheat blast pathogen, but the absence of strict host specialization among the major sub-clades suggests that the ability to infect wheat may have originated multiple times. All of our findings are consistent with the hypothesis that wheat blast emerged in Brazil through a host shift from the *Pygt* population infecting other grasses growing near wheat fields, with strong evidence that gene flow still occurs between the *Pygt* population infecting wheat and the *Pygt* population infecting other grasses. We hypothesize that this recurring gene flow enables *Pygt* populations to maintain significant genetic variation on multiple hosts, a finding that stands in stark contrast to what is found for populations of *P. oryzae* causing rice blast.

The microsatellite and virulence datasets revealed that the contemporary *Pygt* population of Brazil possesses a high degree of genetic and phenotypic diversity. We identified 198 MLMGs and 25 virulence groups among 526 *Pygt* isolates.

We found little differentiation among populations infecting wheat and other grasses, indicating that *Pygt* is not a wheat-specialized pathogen. Populations separated by more than 2000 km were very similar, indicating a high degree of gene flow across large spatial scales and/or high levels of genetic diversity, which would reduce the impact of genetic drift and maintain similar allele frequencies over longer periods. The high gene flow may reflect efficient wind-dispersal of conidia and/or ascospores as well as long distance dispersal on infected seed of wheat and *Urochloa* [72]. Gametic equilibrium was found among SSR markers in most populations, with both mating types present, though with a predominance of the *Mat1-1* idiomorph. These findings, coupled with both high genotype diversity (198 MLMGs out of 526 total strains analyzed) and evidence for some clonality, indicate that *Pygt* has a mixed reproductive system in which cycles of sexual reproduction are followed by the dispersal of locally-adapted clones. The absence of shared MLMGs between populations sampled in 2005 and 2012 suggest that clones do not persist for long periods of time, unlike what has been reported for *P. oryzae* [73]. Alternatively, very high genetic diversity would make it less likely to find the same MLMGs among populations.

Historical analyses of gene flow indicated significant genetic exchange between *Pygt* populations on wheat and other grasses, with the direction of gene flow predominantly from the population infecting other grasses and into the populations infecting wheat. We hypothesize that the fungal strains capable of infecting both wheat and other grasses can move back and forth between hosts, with recombination occurring mainly on the other grasses and giving rise to the highly diverse *Pygt* population we observe today. Support for this scenario can be found in previous reports of cross infection and inter-fertility between isolates from wheat and other poaceous hosts [52–54], as well as in the lack of differentiation among wheat- and other Poaceae-adapted populations, the sharing of genotypes and virulence groups between the two host groups, and the finding of gametic equilibrium consistent with sexual recombination in most populations.

The finding that populations of *Pygt* from wheat and other grasses were not genetically subdivided suggests that several grass species can be hosts for the wheat blast pathogen, unlike the strict host specialization observed for the rice blast pathogen. We hypothesize that *Pygt* spends most of its life cycle colonizing grass species neighboring or invading the wheat fields affected by wheat blast. We further postulate that sexual recombination takes place mainly or exclusively in these other poaceous hosts, generating most of the genetic diversity observed in the *Pygt* populations infecting wheat. Other crop pathogens, especially rusts, are also known to undergo sexual recombination on a non-crop host. These hypotheses are consistent with earlier observations that the forage species signal grass (*U. brizantha*) plays a major role in the genetic variation of the wheat blast pathogen by providing a niche for the fungus to sexually reproduce [15, 54]. Because *U. brizantha* is a widely grown forage grass occupying more than 90 million ha in Brazil [74], and is often found growing next to wheat fields, we propose that *U. brizantha* constitutes a major reservoir of wheat blast inoculum and provides a temporal and spatial bridge that connects wheat crops between growing seasons and across the wheat growing areas of Brazil.

Virulence phenotyping of 173 *Pygt* strains differentiated 25 seedling- (SVG) and nine head-virulence groups (HVG). Many wheat cultivars that are resistant to leaf infections are susceptible to head infections, in agreement with the earlier findings [1]. SVG A and HVG A were capable of causing blast on the entire set of tested cultivars. The isolates in these virulence groups form a “super race” that occurs at a relatively high frequency on Brazilian wheat and are also found on *Avena sativa* (N = 10), *U. brizantha* (8), *Chloris distichophylla* (4), *Echinochloa crusgalli* (4), *Rhynchelytrum repens* (4), *Digitaria sanguinalis* (3), *Eleusine indica* (2), *Eragrostis plana* (2), *Cenchrus echinatus, Cynodon* spp., *D. insularis*, *Panicum maximum*, and *S. sudanense*.

The closely related rice blast pathogen *P. oryzae* is often presented as a model for understanding wheat blast. *P. oryzae* populations are considered strictly asexual [75], except for rare sexual populations of *P. oryzae* associated with rice in South-eastern Asia (the origin of rice domestication, and the proposed center of origin for rice blast) [73, 76], and the population associated with finger millet (*Eleusine coracana*) in West Africa. The *Pyricularia* population adapted to finger millet is probably a new *Pyricularia* species distinct from *P. oryzae*, with a center of origin in western Kenya and north-eastern Uganda [77]. However, it is yet to be reclassified. Remarkably, sexual perithecia have not been found in the field for either of these sexual populations, illustrating the challenge of proving a population is sexual even when it exhibits the population genetic "signature of sex" composed of gametic equilibrium among neutral markers, low clonality and mating types at equal frequencies. As was the case for the sexual *Pyricularia* populations on rice in Southeast Asia and on finger millet in West Africa, we have not yet found natural perithecia of *Pygt* in Brazilian wheat fields, but we have abundant population genetic and biological evidence that strongly indicate the occurrence of sexual *Pygt* populations in Brazil.

Our biological evidence for sexual reproduction is the formation of proto-perithecia and perithecia of *Pygt* on autoclaved wheat stems and on senescing stems of wheat and other grasses. Moreira [78] conducted similar experiments by injecting stems of living wheat plants with the same sexually compatible isolates. In that experiment, no sexual structures were produced in living plant tissues [78]. These contrasting results suggest that senescent plant tissues are necessary to stimulate sexual reproduction in *Pygt*. The same pattern emerged when sexually compatible isolates of *P. oryzae* were placed on living rice plants: perithecia formation occurred only in senescent or detached leaf sheaths [79].

While perithecia produced in our assays did not harbor detectable asci and ascospores, the induction of sexual structures in Ascomycetes is known to be affected by many factors including substrate, light, temperature, and the availability of female fertile strains [80]. We hypothesize that the lack of ascospore production in our assays results from one or more of these factors. We suggest that future studies aiming to identify perithecia of *Pygt* in the field should focus on poaceous hosts such as *Phalaris canariensis* that support the development of fully formed perithecia.

Based on all of the existing knowledge of *Pygt* biology and epidemiology, we propose a provisional disease cycle for wheat blast (Fig 10). At the end of a cropping season (Ae), ear infections lead to infected seed (B, C), providing inoculum for both local and long distance dispersal of the pathogen [72]. Crop residues left in the field after harvest provide a niche for *Pygt* sexual reproduction (D, 1-4); the resulting perithecia release airborne ascospores (D1) that create new genotypes that can cause new infections locally or in distant host populations by the germination of terminal cells (D2), which is followed by fungal vegetative growth and subsequent conidiogenesis (D3) [81]. The asexual conidia produced in the resulting infection are released (D4) and provide airborne inoculum for leaf infection on other grasses located within or next to wheat fields (E, F) [1, 53, 82]. Perithecia can also form in other infected poaceous hosts and on major pasture grasses, with the resulting ascospores falling onto nearby wheat crops (E). Seedborne inoculum (B, C) results in primary infections in newly established wheat crops. (F4) conidia released from leaf blast lesions on other poaceous hosts growing near wheat fields can also contribute inoculum leading to blast on wheat ears [1, 53]. Conidia production on leaves (Af) in the lower canopy of some wheat cultivars can coincide with spike emergence in the field and provide an important source of inoculum for wheat blast epidemics on ears (Ae) [83].

**Fig 10.**
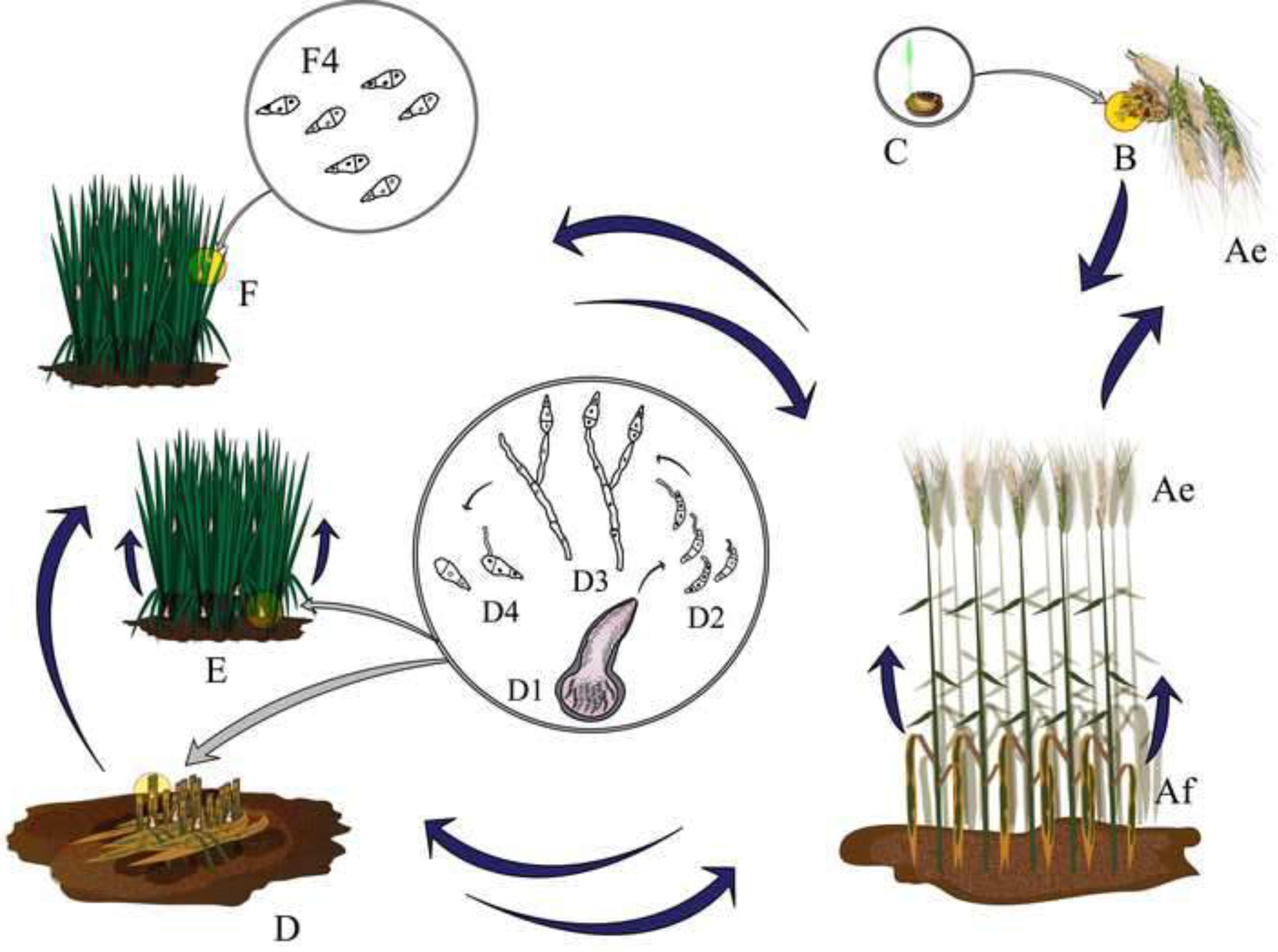
*Pyricularia graminis-tritici* life cycle and wheat blast disease cycle. At the end of a cropping season (Ae), wheat blast infection on ears will result in seed infection (B, C), providing inoculum for either local or long distance dispersal of the pathogen [72]. Crop residues remaining in the field after harvesting, especially under no tillage conditions, serves as a niche for sexual reproduction of the fungus (D, 1-4); the resulting mature perithecia release ascospores by deliquescence of asci (D1), giving rise to new fungal individuals by the germination of terminal cells (D2), which is followed by fungal vegetative growth and subsequent conidiogenesis (D3) [81]; primary conidia originating from this process are released (D4) and constitute airborne inoculum for leaf infection on other poaceous hosts, either invasive or contiguous to wheat fields (E, F) [1, 53, 82]. Perithecia can be formed also in other infected poaceous hosts and major pasture grasses and ascospores released out onto a nearby wheat crop (E). Seedborne inoculum (B, C) results in primary infections in a newly established wheat crops. (F4) conidia released from leaf blast lesions on other poaceous hosts nearby wheat crops also contributes inoculum for wheat blast on ears [1, 53]. Conidia production on leaves (Af) in the lower canopy of certain wheat cultivars coinciding with spike emergence under field conditions and could be an important trigger for wheat blast epidemics on ears (Ae) [83].

In summary, our experiments showed that Brazilian *Pygt* populations maintain very high levels of genetic diversity and are able to infect a surprisingly wide array of grass hosts. *Pygt* populations exhibit a mixed reproductive system and are characterized by high levels of gene flow over long distances. There is evidence for substantial genetic exchange between *Pygt* populations infecting wheat and *Pygt* populations infecting nearby grasses. This combination of properties is likely to make wheat blast a particularly difficult disease to control. We hypothesize that the majority of sexual recombination is occurring on nearby poaceous hosts and that *Urochloa brizantha*, as the major pasture grass in Brazil, plays an important role as a host that provides a steady source of inoculum that connects wheat crops across Brazil.

## Material and methods

### Population sampling

A total of 556 isolates of *Pyricularia* spp. were characterized in this study, comprising ten regional populations sampled from wheat or other poaceous hosts. 526 of these isolates were found to be *Pygt* while 30 isolates were found to be different *Pyricularia* species. Six populations of *Pygt* (387 isolates) were collected from symptomatic heads during the 2012 and 2013 cropping seasons in naturally infected wheat fields in Rio Grande do Sul (RSw), Paraná (PRw), Mato Grosso do Sul (MSw), São Paulo (SPw), Minas Gerais (MGw), Goiás and the Federal District (DF-GOw). The isolates from Distrito Federal and Goiás were grouped into a single population because these locations comprise a single cropping region. *Pygt* strains from wheat fields were sampled along transects as described previously [26]. A seventh *Pygt* population was composed of 79 isolates with distinct multilocus SSR genotypes representing the *Pygt* diversity found in the major Brazilian wheat-growing areas in 2005 [39] (Table 1, Supplementary Table 1).

Two additional *Pygt* populations comprised isolates sampled from other poaceous hosts commonly growing as invasive grasses or weeds located within or nearby wheat fields. The two populations from other poaceous hosts (60 isolates) were sampled from within or nearby three wheat fields in Londrina County, Paraná state (PR_P_), and six wheat fields in Dourados County in Mato Grosso do Sul state (MS_P_). For each field, infected leaves were sampled from invasive grass species exhibiting typical blast symptoms located either within the wheat field or less than 100 m from the edge of the wheat field. The Poaceae species sampled included: *Avena sativa, Cenchrus echinatus, Chloris distichophylla, Cynodon* spp., *Digitaria insularis, Digitaria sanguinalis, Echinochloa crusgalli, Eleusine indica, Eragrostis plana, Panicum maximum, Rhynchelytrum repens, Sorghum sudanense, and Urochloa brizantha*.

### Inference of genealogical relationships among *Pyricularia graminis-tritici* and other *Pyricularia* species

We performed population genomics analyses using single nucleotide polymorphisms (SNPs) across the genome. For the population genomic analyses, the samples included 47 rice blast-associated *P. oryzae* strains with publically available genome sequences, 32 Brazilian strains of *P. graminis-tritici* sampled from wheat and other poaceous hosts, two isolates of *P. oryzae* from *Hordeum vulgare*, two isolates of *P. grisea* from *Digitaria sanguinalis*, two isolates of *Pyricularia* spp. from *Setaria italica* and *Eleusine indica*, one isolate resulting from a cross between K76-79 (from weeping lovegrass, *Eragrostis curvula*) and WGG-FA40 (from finger millet, *Eleusine coracana*) and four wheat blast transcriptome samples collected in Bangladesh in spring 2016 [31]. Among the 32 Brazilian *Pygt* strains sampled between 2005 and 2013, 22 were wheat-infecting strains included in an earlier analysis to infer the origin of wheat blast in Bangladesh [31] and 10 were new blast strains sampled from other grasses and included in this paper. Transcriptomic (RNA) SNPs were identified based on short read alignments against the *P. oryzae* reference genome 70-15, available at Ensembl Fungi (http://fungi.ensembl.org/Magnaporthe_oryzae/Info/Index). For all the completely sequenced genomes, we used Bowtie version 2.2.6 [84] to align quality-trimmed Illumina short read data against the reference genome 70-15. Quality-trimmed Illumina short read data generated from RNA from the Bangladesh transcriptomic samples were mapped using TopHat version 2.0.14 [85]. The variants in the genomes of the different strains were identified using the Genome Analysis Toolkit (GATK) version 3.5 available at the Broad Institute (https://software.broadinstitute.org/gatk/) [86]. A two-step variant calling was used following the GATK best practice guidelines. Firstly, raw variants with local reassembly of read data were called using Haplotype Caller. All the raw variant calls and filtration were jointly genotyped using the GATK Genotype GVCFs. Secondly, SelectVariants was used to subset the variant calls to contain only SNPs. Finally, we applied SNPs hard-filters to remove low-quality SNPs using the following criteria: QUAL ≥ 5000.0, QD ≥ 5.0, MQ ≥ 20.0, − 2.0 ≤ ReadPosRankSum ≤ 2.0, −2.0 ≤ MQRankSum_upper ≤ 2.0, −2.0 ≤ BaseQRankSum ≤ 2.0. Furthermore, we used vcftools (https://vcftools.github.io) to generate a SNP dataset for phylogenomic analyses. To avoid biases in the phylogenetic reconstruction, we only retained SNPs that were called in at least 90% of all analyzed strains. Furthermore, we retained a SNP only if the SNP was called in the best-sequenced Bangladesh sample 12, as described previously [31] (Supplementary Table S1). We retained 55,041 informative SNPs. A maximum likelihood phylogeny was constructed from a SNP supermatrix using RAxML version 8.2.8 (http://www.exelixis-lab.org) with a GTR substitution matrix and 100 bootstrap replicates.

### Microsatellite genotyping and fragment analyses

526 *Pyricularia* isolates (Table 1, Supplementary table 1) were genotyped for 11 microsatellite loci (cnpt_mg-c013Tri, -c047, -c060, -c065, -c108, -c129, -c147, -c168, -c233, -c248, and -p1e11) as described earlier [39, 87] (Supplementary Table 2). Briefly, amplifications were performed in a thermal cycler with conditions as follows: initial denaturation at 95°C for 3 min; followed by 35 cycles of 95°C for 25 s, 55°C or 60°C for 25 s, and 72°C for 25 s; with a final extension of 72°C for 15 min. PCR reactions were diluted and combined in three sets for electrophoresis (Set 1: cnpt-mg-c047, -c065, -c248, and -p1e11; Set 2: cnpt-mg-c013Tri, -c060, -c147, and -c168; and Set 3: cnpt-mg-c108, -c129, and -c233). Isolates 12.1.111 and 10880 were included as controls in every run of 93 samples. The fluorescent-labeled PCR products, along with a size standard were separated on an ABI 3730xl capillary sequencer. The fragment analysis for detection and discrimination among allele sizes was performed using Geneious R 9.1.5.

### Analyses of population genetic structure

SSR datasets were used to calculate gene and genotype diversity and genetic differentiation among populations, generate minimum spanning networks among genotypes, and estimate contemporary patterns of migration and gene flow. We inferred the predominant reproductive mode based on tests of gametic equilibrium and frequencies of the mating type idiomorphs *Mat1-1* and *Mat1-2*. Except for the analyses of genotypic diversity, all analyses used clone-corrected datasets in which only one individual from each multilocus microsatellite genotype was included per population.

### Genotypic and genetic diversity and allelic richness

The multilocus microsatellite genotype (MLMG) for each isolate was determined using Genodive v. 2.0b7 [57]. Isolates exhibiting the same MLMG were considered clones. A minimum spanning network (MSN) was constructed to show the distribution and genetic similarity among the MLMGs of *Pygt* found in the nine populations. The MSN was constructed with the *bruvo*.*msn* distance function [88] and the *Prim* algorithm of the *igraph* package, [89] using the *poppr* package [90] in the R environment [91].

Measures of genotypic diversity included: a) number of MLMGs per population; b) population-specific MLMGs; c) clonal fraction calculated as 1-(number of MLMGs)/(total number of isolates); d) effective number of MLMGs (*Go*) [92]; and e) the evenness, an indicator for how evenly the genotypes were distributed in the population, calculated as the ratio of the effective number of distinct MLMGs scaled by the maximum number of expected MLMGs. We tested the statistical significance of differences in genotypic diversity between pairs of populations based on 1,000 bootstrap resamplings matching the size of the smallest population (19 individuals) [61]. Allelic richness was estimated for each population as the average number of alleles per locus using rarefaction [58]. To test whether populations differed in allelic richness, *p* values for the significance of the pairwise comparisons were obtained by 1,000 permutations. These calculations were computed using FSTAT v. 2.9.3.2 [93]. The probability of identical genotypes arising from sexual reproduction and random mating and it is identical to the genotype probability was estimated with the *Pgen* index previously described with GenAlEx v6.501 software [55, 56]

### Population differentiation

AMOVA [94] was used to assess the distribution of gene diversity and the degree of differentiation among geographical populations of the pathogen. Populations were also grouped according to the host of origin. Degrees of differentiation were compared using orthogonal contrasts. The sum of squared size differences (*R*_*ST*_) was used as the distance measure between two haplotypes [60]. The significance of the fixation indexes was tested using 1,023 permutations by a nonparametric approach [94] at α = 0.05 after Bonferroni correction for multiple comparisons [62]. All calculations were carried out with the program ARLEQUIN v. 3.11 [59].

### Assessment of historical migration and demographic parameters

For migration analyses, populations were grouped according to their host of origin. A maximum likelihood test based on MCMC [68] was used to test four different models of migration between the populations obtained from wheat and from other Poaceae. The migration models tested were: a) complete panmixia; b) bidirectional; c) directional, with migration occurring from the wheat population towards the other Poaceae population; and d) directional (inverse) with migration occurring from the population obtained from other Poaceae towards the wheat population. Estimates of gene flow were obtained using five runs, and the run with the highest likelihood was chosen to represent each migration model. Then the likelihood values of the four migration models were compared to select the one that best fit the data based on the Log of the Bayes Factor (LBF). LBF was calculated as 2 [ln(Prob(Data | ModelX)) − ln (Prob(Data | best of the four models))]; higher LBF values reflect better fits of the migration model to the data [64, 66].

For all migration analyses the data type chosen was microsatellite data with Brownian motion and assuming a stepwise mutation model. Each of the five runs had ten short initial chains, one long final chain, a static heating scheme with five temperatures (1, 100, 1000, 10,000 and 100,000), and swapping interval of 1. The initial chains were performed with 500-recorded steps, a sampling increment of 100, with 2,500 trees recorded per short sample. The long chain was carried out with 8,334-recorded steps, a sampling increment of 500, six concurrent chains (replicates) and 500 discarded trees per chain (burn-in). The final number of sampled parameter values was 25,002,000 iterations. The values and confidence intervals for the migration rate (*M*), and the effective population size (θ = 2Neμ for haploids, where Ne = effective population size and μ = mutation rate inferred for each locus) were calculated using a percentile approach. Migration analyses were implemented in MIGRATE-n v. 3.6.11 [64] at the CIPRES Science Gateway [63].

### Tests for gametic equilibrium

Gametic equilibrium was assessed using a multilocus association test (10). The hypothesis that genotypes at one locus are independent from genotypes at another locus was tested using Fisher’s exact test at α = 0.05 and an MCMC algorithm (with 1,000 batches and 1,000 iterations/batch) implemented using the program GENEPOP v.3.4 [69]. The Bonferroni correction was applied to this analysis to avoid false rejections of the null hypothesis due to the large number of comparisons performed [62]. Two loci were in gametic equilibrium when their associated *p* value was not significant (*p* > 0.05). We also measured the indexes of multilocus association (*I*_*A*_ and 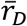) for each *Pygt* population using Multilocus software ver 1.3b, according to Agapow and Burt [70].

### Determination of mating type idiomorphs

The mating type idiomorph, *Mat1-1* or *Mat1-2* [95], was determined for each strain using a PCR assay [39]. To amplify*Mat1-1*, the primers were A1:5’-AGCCTCATCAACGGCAA-3’ and A5: 5’-GGCACGAACATGCGATG-3’. For *Mat1-2* they were B15: 5’-CTCAATCTCCGTAGTAG-3’ and B16: 5’– ACAGCAGTATAGCCTAC-3’. We included isolate Py46.2 as a positive control for *Mat1-1* and a negative control for *Mat1-2*, and isolate Py5003 as a positive control for *Mat1-2* and a negative control for *Mat1-1* [39].

### Development of *Pygt* perithecia on senescing stems from several poaceous hosts

*Pygt* strains Py33.1 (*Mat1-1*) and Py05046 (*Mat1-2*) were shown to be fertile in earlier studies [39, 78]. The production of perithecia and asci on autoclaved wheat stems and naturally senescing stems of wheat and other grasses was assessed after co-inoculation with these strains. The other poaceous hosts assayed were: *Avena strigosa* (black oats) cv.

Embrapa 29 Garoa; *Hordeum vulgare* (barley) cvs. BR Elis and MN 743; *Oryza sativa* cvs. BRS Primavera, BRSMG Relampago and Yin Lu 30 (red rice); *Phalaris canariensis* (canary grass); *Secale cereale* (rye) cv. BR1; *Setaria italica* (foxtail millet); *Triticum aestivum* (wheat) cvs. BRS 264 and MGS Brilhante; Triticale (x*Triticosecale*) cv. IAC Caninde; *Urochloa* hybrid cv. Mulato (*Urochloa ruziziensis* × *U. decumbens* × *U. brizantha*). The wheat cv. MGS Brilhante is classified as moderately resistant to wheat blast, while the other wheat cultivar, barley, *Urochloa* spp., and oats are considered susceptible to wheat blast. In contrast, rice cultivars are resistant to *Pygt* [15, 39]. The remaining hosts included in this experiment have unknown susceptibility to *Pygt*.

Spores of isolates Py33.1 and Py05046 were harvested after 14 days of growth on oatmeal agar [39] and combined in equal proportions at 1×10^4^ conidia ml^−1^ for co-inoculation as described earlier, with minor modifications [79]. Wheat stems consisted of 4-cm sections collected from one-month old plants and autoclaved at 121°C for 20 min. Autoclaved wheat stems or naturally senescing stems were placed in 90 mm Petri dishes containing water agar (agar, 15gl^−1^) and were inoculated by injection of 0.3 mL of the spore mix. Inoculated materials were kept in a growth chamber at 25°C under a 12 h dark /12 h fluorescent white light photoperiod for 7 days. Subsequently, for perithecia development, the temperature was lowered to 200C and the samples were incubated for another 21 days (autoclaved stem sections) or one month (senescing stem pieces) under the same photoperiod. The assays were replicated once, with five repeats of each experimental unit each time. The development of sexual structures was documented using light and scanning electron microscopes. The density of proto-perithecia or perithecia on plant debris and on sections of senescing stems was determined by analyzing at least three areas of approximately 0.5 mm^2^ on each plant species.

### Virulence spectrum of *Pygt* on wheat seedlings and detached heads

The virulence spectra of 173 isolates of *Pygt* representing 80 MLMG were assessed on seedlings and detached heads of ten wheat cultivars and one barley cultivar. Within each MLMG, isolates were selected at random from the eight populations sampled in 2012-2013, including 121 isolates from wheat and 52 isolates from other poaceous hosts. The wheat cultivars included in the tests were: Anahuac 75 (susceptible control), BR 18, BR 24, BRS 220, BRS 229, BRS 234, BRS Buriti, CNT 8, MGS 3 Brilhante, Renan, and barley cv. PFC 2010123.

Detailed procedures for inoculum preparation, inoculation, incubation, disease assessment and data analysis were described earlier [15, 39]. Briefly, inoculations were conducted on 15-day-old seedlings at the 4-leaf stage and on detached heads harvested from plants after anthesis. Seedling and head inoculation experiments were conducted using a two-factor completely randomized balanced design. Two pots containing ten plants each were used for the seedling test, while three foam blocks with ten detached heads apiece were inoculated for each of the 173 isolates. Each inoculation test was conducted twice. In both tests, disease was scored 5 days after inoculation. Cultivars were classified as resistant (R) or susceptible (S) based on visual assessment of the percentage of leaf or detached head showing typical blast symptoms. *Pygt* isolates were placed into seedling virulence groups (SVGs) and head virulence groups (HVGs) according to their pathogenicity spectra on each wheat cultivar.

## Acknowledgements

Primer sequences for MAT loci were provided by Didier Tharreau, INRA, Montpellier, France.

## Supporting information

**S1 Table. Isolates of *Pyricularia* species included in the inference of genealogical relationships.** This table lists and describes all the isolates included in the inference of genealogical relationships among wheat blast samples of *Pyricularia graminis-tritici* and several other blast samples.

**S2 Table. Isolates of *Pyricularia* species analyzed in this study.** This table lists and describes all the isolates examined in this study, including their original host, year and location of sampling, mating type, multilocus microsatellite genotype, alleles found in 11 microsatellites loci, and seedling and head virulence group for each isolate.

**S3 Table. Oligonucleotides.** This table lists all primers for microsatellite loci and their sequences in this study.

## Author Contributions

**Conceptualization:** PCC, JLNM, EA, BAM
**Data Curation:** VLC, ALDD, JTAR, ALVB, CAF, JLNM, SIM, PCC, EA, DC
**Formal Analysis:** VLC, PCC, SIM, DC
**Funding Acquisition:** JLNM, EA, PCC, BAM, DC
**Investigation:** VLC, ALDD, SIM, JTAR, GC, JLNM, PCC, DC
**Methodology:** VLC, ALDD, SIM, GC, ALVB, JLNM, PCC, DC
**Project Administration:** VLC, JLNM, PCC, EA, DC
**Resources:** PCC, JLNM, EA, DC, BAM
**Supervision:** PCC, JLNM, CAF, BAM
**Validation:** VLC, ALDD, ALVB, SIM, JTAR, JLNM, EA, PCC, DC
**Visualization:** GC, PCC
**Writing-Original Draft Preparation:** VLC, SIM, PCC
**Writing-Review & Editing:** VLC, JNM, BAM, SIM, JLNM, PCC, DC

